# Haplotype based testing for a better understanding of the selective architecture

**DOI:** 10.1101/2022.07.18.500395

**Authors:** Haoyu Chen, Marta Pelizzola, Andreas Futschik

## Abstract

The identification of genomic regions affected by selection is one of the most important goals in population genetics. If temporal data are available, allele frequency changes at SNP positions are often used for this purpose. Here we provide a new testing approach that uses haplotype frequencies instead of allele frequencies. With this approach, less multiple testing correction is needed, which leads to tests with higher power, especially when the number of candidate haplotypes is small or moderate. Another advantage is that haplotype frequencies can often be recovered with less noise than SNP frequencies, especially under pool sequencing. For a larger number of haplotypes, we investigate methods to combine them to a moderate number of haplotype subsets. The use of haplotypes also permits a better understanding of selective signatures. For this purpose, we propose post hoc tests for the selected haplotypes and differences between their selection coefficients. Using both simulated and real data sets, we illustrate the performance and benefits of our proposed test statistics.

## 1 Introduction

Evolve and Resequence (E&R) experiments [18] provide a modern approach for studying patterns of adaptation in a controlled environment. In these experiments, one or multiple populations are followed over time, often under stressful environmental conditions. Researchers then aim to identify adaptive changes at a genetic level. High-throughput whole genome sequencing techniques provide allele and haplotype frequency data at suitable time points during the experiment. Depending on the experimental design and the available resources, sequencing can be performed at the beginning and the end of the experiment or at multiple time points. The experiment is often also replicated so that analogous data are obtained from multiple populations.

Once data are available, statistical tests are often used to identify the presence of selection. A proper test needs to take all sources of random variation into account. Indeed, besides selection, observed allele frequencies are affected by genetic drift, and frequently also by sampling and sequencing noise. So far, different SNP based tests for selection have been proposed in the context of E&R experiments. Some approaches, such as [19] are heuristic and do not control the type I error. More recently, in [7] a modified version of the classical chi-square and CMH test has been proposed that is able to take all relevant sources of randomness into account. For a review of further available methods, we refer to [44].

Here we propose tests that rely on haplotype frequencies instead of SNP frequencies and illustrate their potential and advantages. In the different context of genome-wide association studies (GWAS), testing based on haplotypes has already been used [25, 23] to identify genetic variants that are associated with phenotypic traits of interest. Available methods include likelihood ratio tests [22] and score tests [21], and recently a combination of haplotype block and SNP set approaches has been proposed in [24].

For samples from natural populations, yet further examples of haplotype based tests can be found in [45] and its extension [2], where signatures of recent selection are found by searching for regions with long range linkage disequilibrium. The haplotype based test proposed in [46] is similar in spirit to [45] but focuses on ongoing selection.

Given that haplotype based testing has already proved promising in the above mentioned setups, it seems desirable to make it available also for E&R experiments. Since this involves temporal allele frequency data and a different null model, our proposed tests require a new methodological approach that is explained in detail in Section 2. In summary, we identify selected genomic windows by testing each haplotype against the combination of all others using a modification of the chi-square or (with replicate populations) the Cochran-Mantel-Haenszel (CMH) test [7]. The tests take all relevant sources of random variation into account. Subsequently, we combine the resulting p-values using recently proposed combination tests for the global null hypothesis.

When the actual haplotype frequencies are not available, haplotype reconstruction techniques [17, 8] provide the possibility of estimating this information from allele frequency data. The standard error of these estimates will then be one of the sources of random variation.

In our simulations, we observed that haplotype based tests do not necessarily outperform SNP-based methods if the total number of haplotypes gets too large. Therefore we propose two variants of our approach in subsection 2.4 that provide improved power with experiments involving many haplotypes.

Furthermore, when the presence of selection is established by the haplotype based test, we provide a post hoc test for the number of selected haplotypes in subsection 2.2. In subsection 2.3, we propose another post hoc test for differences in fitness between pairs of haplotypes. We provide results on simulated data for these two tests and show that they have good power under many scenarios (Section 3). In Section 4, we also apply our proposed tests on real data considered by [35].

## 2 Methods

Consider a haploid population with effective population size *N*_*e*_, and a genomic region exhibiting *N*_*Hap*_ haplotypes. We summarise the temporal dynamics of these haplotypes over *T* + 1 generations via the relative haplotype frequencies 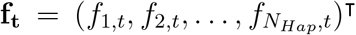 at generation *t* (0 ≤ *t* ≤ *T*). Without selection, the relative haplotype frequencies in a subsequent generation are obtained via multinomial sampling from the previous generation [1]:

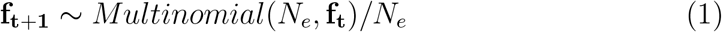

The changes in haplotype frequencies caused by repeated multinomial sampling are commonly known as genetic drift. Under selection, the haplotypes differ in fitness, leading to modified multinomial sampling probabilities [1]:

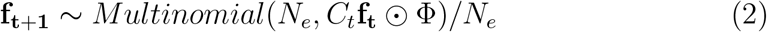

where

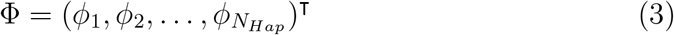

is the fitness vector, *C*_*t*_ a normalising constant at time *t* such that 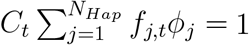, and ⊙ the Hadamard product. Neutrality corresponds to the special case where all elements of Φ are equal and 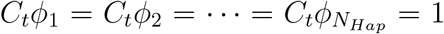. Otherwise, higher fitness of a certain haplotype compared to others represents the presence of a selective advantage.

In the context of evolve and resequence, the experiment is often replicated, which leads to R independent haplotype frequency vectors **f**_**t**_^(1)^, **f**_**t**_^(2)^, …, **f**_**t**_^(*R*)^ at any sequenced time point *t*. Suppose we have an experiment with *k* sequenced time points (*t*_0_, *t*_1_, …, *t*_*k−*2_, *t*_*k−*1_), with *t*_0_ = 0 and *t*_*k−*1_ = *T*. Table 1 displays the haplotype frequency matrix **F**^(*r*)^, for some replicate population *r*. These matrices may then be combined in to a *N*_*Hap*_ *× kR* matrix **F =** [**F**^(1)^ **F**^(2)^ … **F**^(*R*)^]. If the true frequencies are unknown, we use estimates 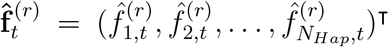, and 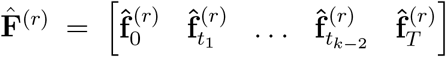 instead of the actual quantities. These estimates will typically contain sampling and sequencing noise that needs to be taken into account when testing hypotheses.

**Table 1:**
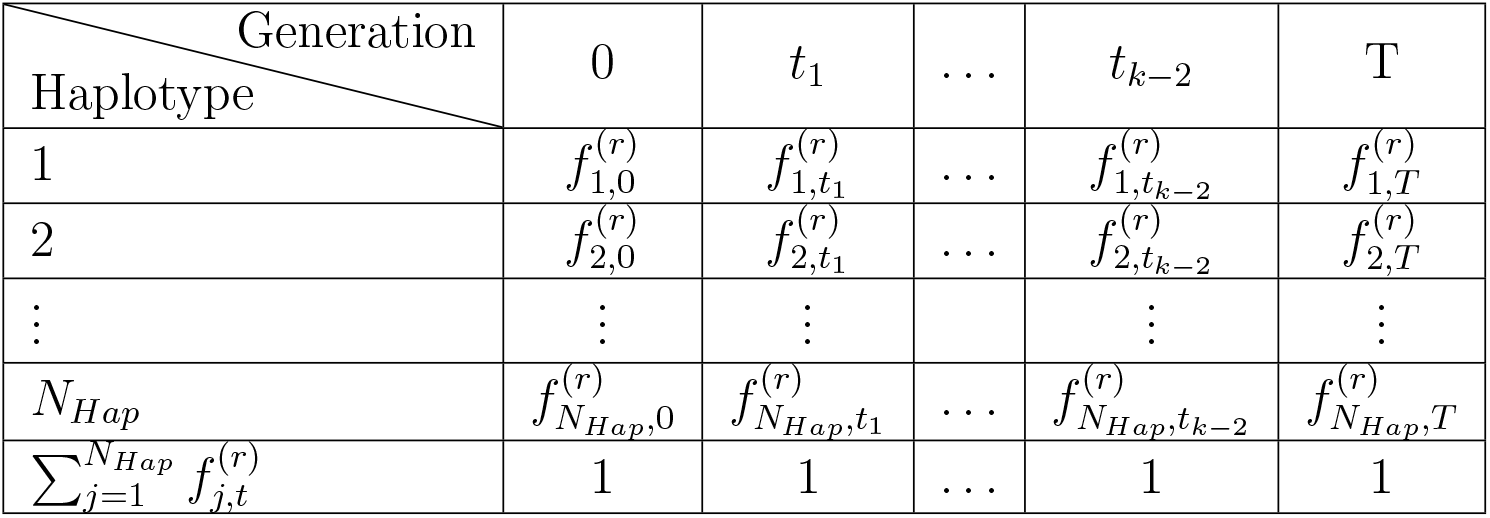
Structure of (true) haplotype frequency matrix with *k* sequenced time points for some replicate population *r*. The estimated frequency of haplotype *j* at generation *t* for replicate *r* is denoted by 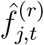.

To test for selection against the neutral null hypothesis 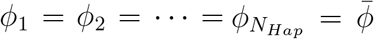 where 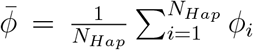 is the mean fitness, we decompose the global null hypothesis into a multiple testing problem. For each haplotype *j* (1≤ *j* ≤ *N*_*Hap*_), distinguishing neutrality from selection may be phrased in terms of the hypothesis testing problem:

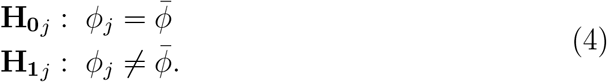

We propose the usage of the adapted CMH test [7] when multiple independent replicate populations are available. It naturally reduces to the adapted chi-square test if there exists only one replicate. The test is conducted in a binary fashion such that for some haplotype *j*, the test is conducted between the frequencies of haplotype *j* across all replicates 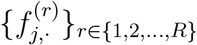 and the cumulative frequencies of all other haplotypes 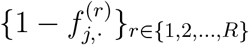.

As discussed in [7] in the context of SNPs, the estimated haplotype frequencies 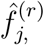 may involve multiple components of variance. For simplicity, we present the test statistic assuming that all haplotype frequencies are known, and the only relevant variance component is genetic drift. For cases where other sources of variance such as sampling and pool sequencing noise are present, the test statistic can be found in Section S.2 (the prefix S-refers to sections/figures/tables in the Supplementary Material). For some haplotype *j*, the adapted CMH test using known haplotype frequencies has the following test statistic:

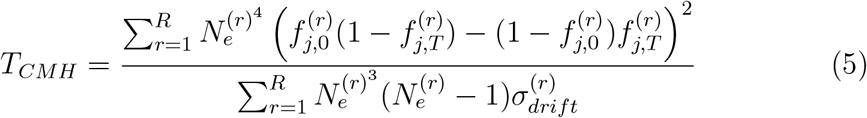

where 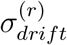 is the variance of haplotype frequencies due to drift at replicate *r*:

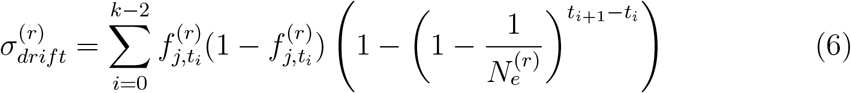

with *k* being the total number of sequenced time points, *t*_0_ = 0, *t*_*k−*1_ = *T* being the first and last time points respectively as before. In practice, the effective population size *N*_*e*_ will often be unknown and needs to be estimated for instance by using the method proposed by [36]. In the special case where no information at intermediate time points is available we have *k* = 2, as only the start and end time points are sequenced. The test is carried out for all null hypotheses *H*_0*j*_, *j ∈* {1, 2, …, *N*_*Hap*_} provided in (4). This leads to *N*_*Hap*_ p-values that are combined by a suitable multiple testing procedure.

Algorithm 1 provides pseudocode that summarizes our approach. Details of the proposed multiple testing approach can be found in subsection 2.1 below.

### 2.1 Multiple testing procedure

We carry out one hypothesis test for each hypothesis pair in (4) and want to test the global null hypothesis (i.e. the null hypothesis for all *j ∈* {1, 2, …, *N*_*Hap*_}). To control the type I error, we need a proper multiple hypothesis testing procedure. In principle, Bonferroni [4] tests or the recently proposed approach by [27] outlined in Section S.1 might be used. However, we found these methods to be too conservative in most situations. Therefore, we focus on more powerful approaches such as the omnibus test [9] and the harmonic mean p-value [10]. Although no theory ensures type I error control for these methods under dependence, most of our simulated scenarios did not lead to violations. However, we observed type I error probabilities that slightly exceeded the significance threshold for both tests in the case of one replicate population, a small number of haplotypes, and known haplotype frequencies. See Section 3.8 for details.

#### Algorithm 1: Haplotype based selection testing

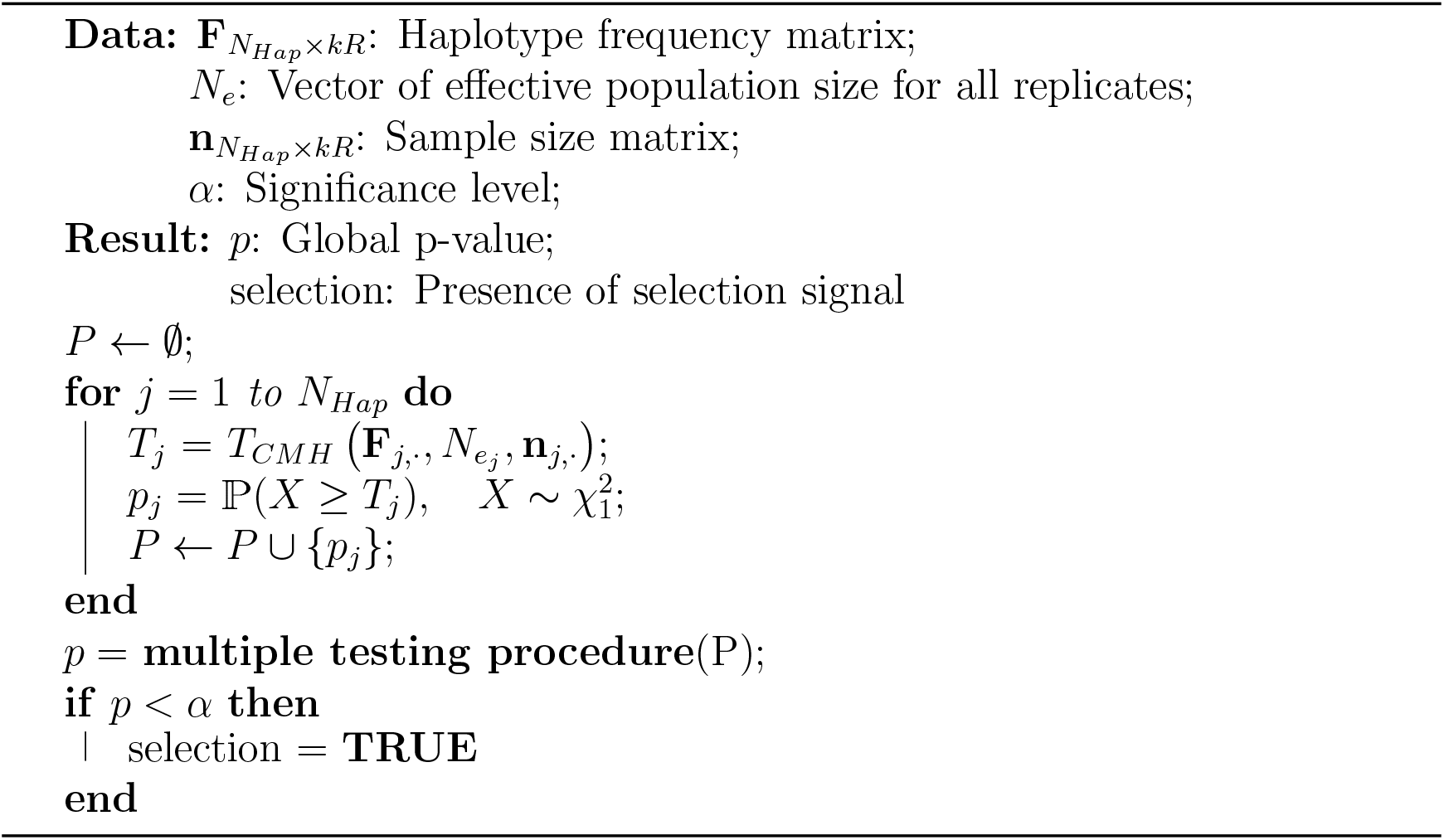

#### 2.1.1 Omnibus test

The omnibus test proposed by [9] has originally been derived for independent p-values, and was shown to provide good power under various deviations from the global null hypothesis. For sorted p-values {*p*_(*j*)_}_*j∈*{1,2,…,*N*}_ such that:

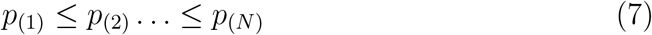

the L-statistic *S*_*i*_ is computed by adding up transformed p-values up to rank *i*. Here we use the proposed default transformation (negative logarithm) leading to

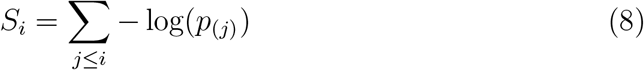

with the test statistic *T* being:

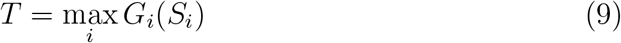

and *G*_*i*_ the cumulative distribution of *S*_*i*_ under the global null hypothesis of neutrality. Note that the assumption of independent p-values is not fulfilled in our context, as the haplotype frequencies at any given time point add up to 1. We observed violations in terms of type I error control only under a few scenarios, see Section 3.3 for details.

#### 2.1.2 Harmonic mean p-value (HMP)

Another combination method, the harmonic mean p-value proposed in [10] is given by

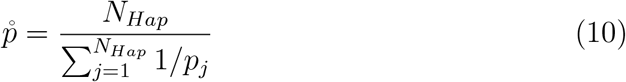

for equal weights. The combined p-value *p* is then calculated as

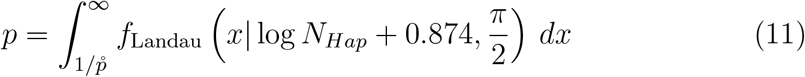

with *f*_Landau_ being the density function of a Landau distribution.

While [39] showed that the type I error is not controlled under some dependence structures, we did not observe any large violations under our considered scenarios. See Section 3.8 for further details.

### 2.2 Testing for the number of selected haplotypes

In association studies and genomic prediction, regression models are often used to explore the influence of haplotypes on a phenotype [23], [48]. While such a response variable is lacking in our setup, further information about the number of selected haplotypes is of interest in our context. Therefore we propose a follow up test for this purpose. As with forward selection methods in regression models, our approach proceeds in a stepwise fashion. At each step a test is carried out at level *α*, and the procedure stops once no more rejection is necessary. We call a haplotype to be positively selected, if there is at least one other haplotype with lower fitness.

A rejection of the global hypothesis (4) implies that there is at least one selected haplotype. To investigate whether there are further selected haplotypes, we identify the haplotype *m*_1_ that provides the maximum change in frequency over time, normalised by variance and gives the largest contribution to the rejection of the hypothesis:

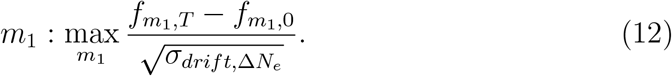

We use

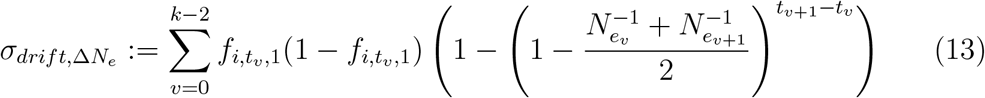

in situations when the true haplotype frequencies are known. This drift variance estimate is more complex than (6). However, it will simply reduce to *σ*_*drift*_ in cases where *N*_*e*_ is constant s.t. 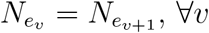. For scenarios where haplotype frequencies are estimated, this variance term will need to change accordingly, see Section S.2.5 for more details.

If 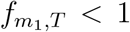, we test whether there are further selected haplotypes. For this purpose, we remove haplotype *m*_1_ and test for fitness differences among the remaining haplotypes:

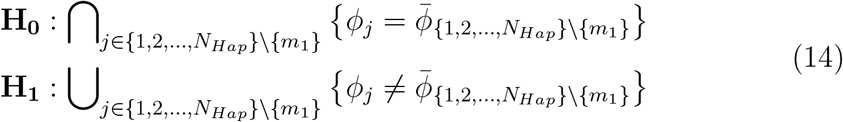

where 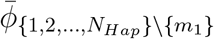 is the mean fitness of all haplotypes except *m*_1_. We furthermore renormalise the remaining haplotype frequencies to add up to 1 at any time point:

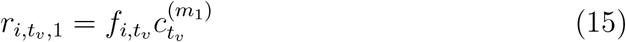

where 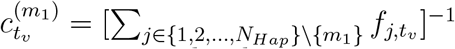. We also recompute *N*_*e*_ separately for each time point to take the removal of haplotype *m*_1_ into account:

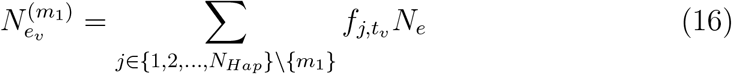

We then test for selection, but replace the variance caused by drift, *σ*_*drift*_ in *T*_*CMH*_ by 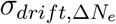. If the null hypothesis is rejected, we claim that there are at least two selected haplotypes in the population.

The above method is then iterated to test for further selected haplotypes. For this purpose, we find *m*_2_, *m*_3_, … respectively by ranking the normalised differences in frequency change over time and test the hypothesis:

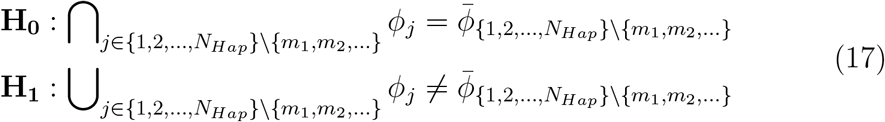

as long as the previous null hypothesis is rejected. The method of normalisation, the *N*_*e*_ computation, and the testing procedure are analogous as before.

If replicates are present, the largest change in haplotype frequency might not be consistent across all replicates. We therefore propose the following criterion to remove haplotypes:

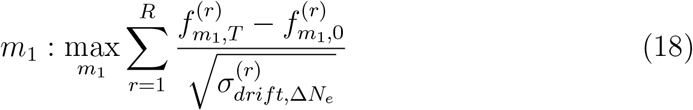

Further haplotypes are excluded in an analogous way. The values of 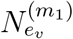 and the frequency normalisation will then be calculated separately for each replicate.

### 2.3 Pairwise test for different fitness across haplotypes

As further post hoc tests, we consider pairwise comparisons for differences in the fitness between haplotypes *i* and *j*:

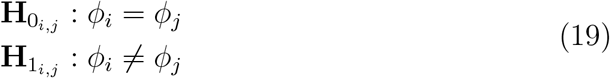

We test this hypothesis pair, if their frequencies satisfy 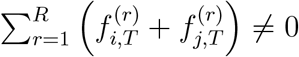.Given the haplotype frequency matrix **F**, for some pair of haplotypes *i* and *j* at replicate *r*, we normalise their frequencies to add up to one. For *l* ∈ {*i, j*}, we set 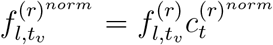, where 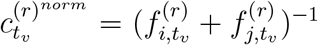 is the normalising constant of replicate *r* at generation *t*_*v*_. Furthermore 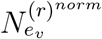 at time point *t*_*v*_ is computed as

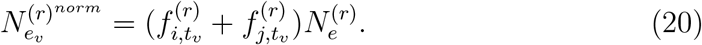

Since this will usually cause a changing *N*_*e*_, we replace the drift variance by 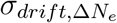 when applying our proposed test (5) to the haplotype pair. To control the false discovery rate, a Benjamini & Hochberg multiple testing correction will also be applied to the p-values obtained from all considered pairs.

### 2.4 Testing when many haplotypes are present

Scenarios with many haplotypes tend to lead to low power when using haplotype based testing. This is due to a large number of individual tests and the small haplotype frequencies. One way to resolve this issue is through the combination of haplotype frequencies. Intuitively, this can be achieved through the removal of SNPs, such that several haplotypes will become identical. Here, we propose two haplotype combination methods to improve the performance of our haplotype based tests.

First, we propose an intuitive approach that combines haplotypes using individual SNP based tests. We use the p-values of these tests without multiple testing corrections and retain SNPs with p-values below some given threshold *β*. The haplotypes that become identical after SNP removal are then combined. A similar approach has been proposed in [42] to identify a selected haplotype. A detailed explanation of this approach is provided in Section S.3.

Another possible approach relies on haplotype blocks obtained via techniques commonly used in GWAS [24]. A haplotype block may be defined as a contiguous region of SNPs that are in high linkage disequilibrium with each other with little evidence of recombination within the region [28]. We use a normalised version of the coefficient of linkage disequilibrium proposed by [29] and follow the approach by [28] to determine haplotype blocks. Our proposed haplotype based selection test is then applied to each of the combined haplotype block frequency matrices. An extra layer of between blocks multiple testing corrections is then needed. We refer to Section S.4 for more details.

## 3 Simulation Experiments

In this section we present the results of an extensive simulation study where we analyse the performance of our proposed tests from Section 2 under different scenarios. First, Section 3.2 provides a proof of concept, illustrating some potential advantages of our proposed haplotype based test compared to SNP based testing for selection in a typical experimental evolution scenario. Then we consider how the choices of the experimental design (Section 3.3) and of the model organism (Section 3.4) can affect the power of our test compared to a SNP based test. Lastly, Sections 3.5, 3.6, and 3.7 illustrate the results of the extensions of our proposed test discussed in Sections 2.2, 2.3, and 2.4 respectively, and Section 3.8 discusses type I error control of our proposed methods.

### 3.1 Data and Simulation setup

Our simulation studies are inspired by the experimental setups described in [40] and in [35]. For our simulations related to the first setup, we randomly selected *N*_*Hap*_ haplotypes (out of 186) and *N*_*SNP*_ SNPs (out of 500) from this window without replacement. Starting from the chosen founder haplotypes, we simulated evolve and resequence experiments with and without selection by generating multinomial haplotype frequency changes along generations using equation (2).

All frequencies are assumed to be known unless otherwise stated. For haplotype frequencies, we set the starting frequencies at time point 0 to be equal, such that each haplotype has a frequency of 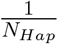. As discussed in Section 4, however, our methods may also be used with arbitrary, non-equal starting frequencies.

Under selection, we consider scenarios involving both one and more than one positively selected SNPs. With *n*_*sel*_ *≥* 1 selected SNPs, we randomly choose positions 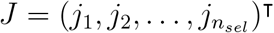 for the selected SNPs. The corresponding vector 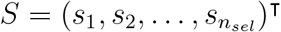 denotes the selection coefficients of these SNPs. Assuming additive fitness effects, this leads to a fitness vector Φ (see equation (3)) with components

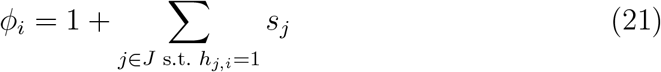

for haplotype *i* (1≤ *i* ≤ *N*_*Hap*_).

We then simulate haplotype frequencies up to (*k* − 1) *×* Δ*t* generations. Given a haplotype structure matrix **H** and a haplotype frequency matrix **F**^(*r*)^ for replicate *r*, the allele frequency matrix **A**^(*r*)^ for this replicate can be calculated as:

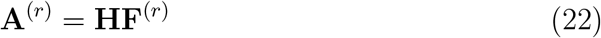

where each element *a*_*i,j*_ denotes the allele frequency for SNP *i* at generation *t*_*j*_. For scenarios where frequencies are estimated with a sample size of 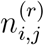, we construct the noisy haplotype frequency matrix 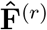 by drawing its columns independently via multinomial sampling. The observed allele frequencies are then obtained as

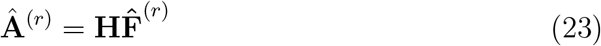

Under pool sequencing with sequencing coverage 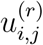 for SNP *i* at time point *t*_*j*_ and replicate *r*, we construct the noisy allele frequency matrix **Ã**^(*r*)^ by drawing each element via binomial sampling using the respective sequencing coverage. With SNP based testing, these binomial variances are added as a component of variance to the denominator of the modified CMH tests (see Section S.2.3).

For the haplotype based test, we assume that the haplotype frequencies are estimated from the noisy allele frequencies **Ã**^(*r*)^ by solving the regression model

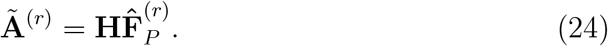

The variances of the estimated regression coefficients 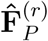 as well as of their sums are then used as additional components of variance.

### 3.2 Proof of concept

As an initial illustration of our hypothesis test we consider 10,000 simulations from a window of 500 SNPs, mimicking an experiment with 10 replicates and a haploid population of size 1,000, with a coverage of 50. Ten founder haplotypes are present in each population. We consider a scenario with 60 generations and with sequencing taking place every 10 generations. To simulate selection, one locus is assumed to be beneficial. In our first example the locus is private to one of the haplotypes and has a selective strength *s* = 0.02. In the second example, the selected locus is shared among five haplotypes and has a selective strength *s* = 0.03. We chose these two selection regimes for illustration purposes, but similar conclusions can also be drawn changing the selection strength.

The receiver operating characteristic (ROC) curves for our proposed haplotype based test and SNP based test under the two scenarios are plotted in Figure 1. We include results under the HMP and the omnibus p-value combination methods.

**Figure 1:**
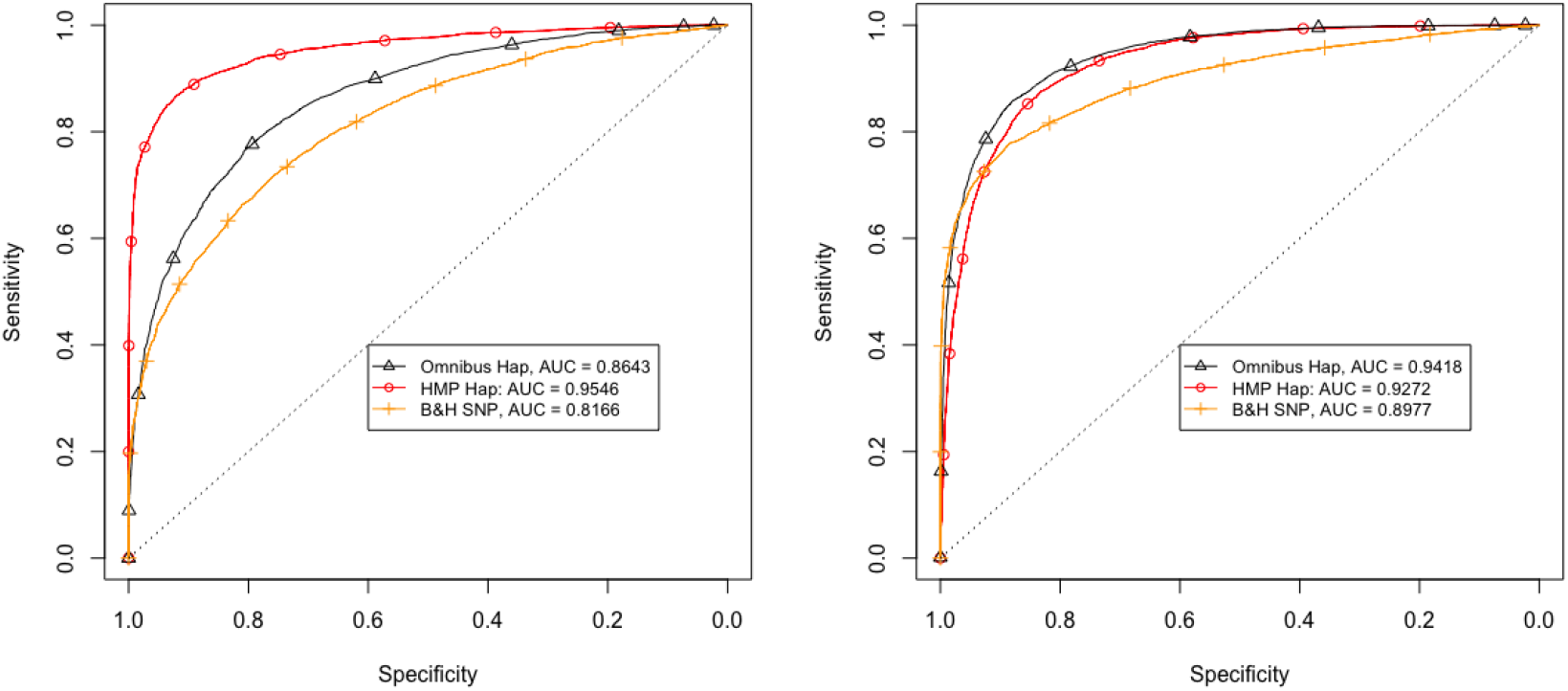
Results for two simulated examples that mimic a typical experiment. The receiver operating characteristic (ROC) curves of the haplotype and SNP based tests are shown for two scenarios: 10 replicate populations with 10 founder haplotypes each are simulated with a coverage of 50. In the left panel, one haplotype is beneficial with *s* = 0.02. In the right panel *s* = 0.03, and 5 haplotypes have a common selective advantage compared to the remaining populations.

The haplotype based tests have a higher area under the curve (AUC) than the SNP based test in both examples. The difference is particularly large under the scenario with one selected haplotype (left panel of Figure 1). This demonstrates that our proposed approach is able to provide a considerable increase in power under some scenarios.

The harmonic mean p-value combination method has particularly high power in scenarios with a single true alternative. Such a situation occurs both when one, or all but one haplotypes are selected. On the other hand, the omnibus test performs better than HMP at intermediate numbers of selected haplotypes (see right panel of Figure 1).

With an intermediate number of selected haplotypes, the advantage compared to SNP based testing is also smaller than for both a large and a small number of selected haplotypes. See Figure S2 for results under scenarios without pool sequencing noise.

### 3.3 Influence of the model parameters

In experimental evolution, different organisms and experimental setups are chosen according to the aims of the experiment and the available resources [47]. Therefore, we discuss the impact of the experimental design on the performance of our haplotype based tests. We investigated the influence of each parameter described in Section 3.1 on the performance of our proposed tests by considering a set of alternative values for each of them. The other parameters have been kept constant as listed in the reference table 2. The results in this section assume known haplotype frequencies unless otherwise stated and are based on the test statistic provided in Section 5.

**Table 2:**
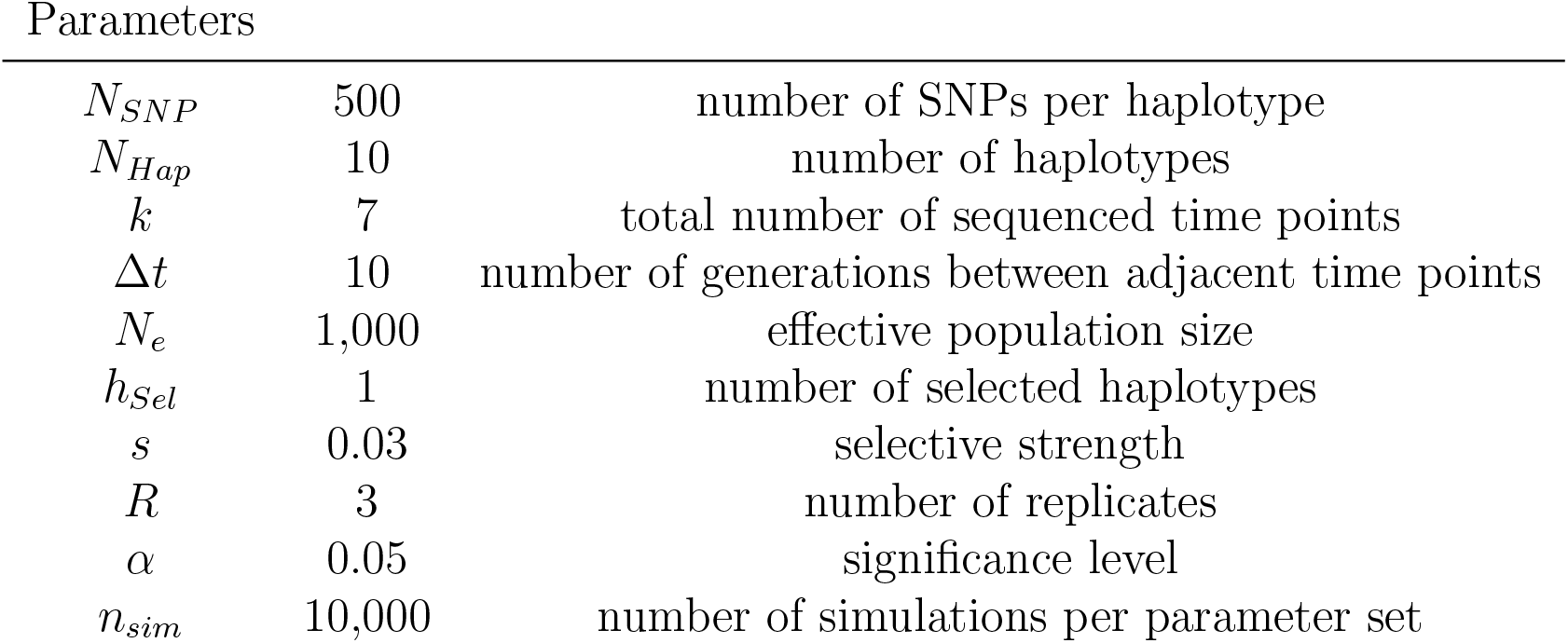
Default parameter values for Sections 3.3, 3.4, 3.5, 3.6, 3.7 and Supplement Section S.6. Unless otherwise mentioned, Figures within these sections use parameters from this section when simulating results.

Figure 2 shows that the power of all tests decreases with an increasing number of initial haplotypes. We observe that the differences in AUC between haplotype and SNP based tests decrease as the number of haplotypes increases. When using the omnibus multiple testing correction, the SNP based approach also slightly outperforms our haplotype based test, if the number of starting haplotypes is large. This can be explained by more multiple testing corrections needed due to the increase in the number of haplotypes.

**Figure 2:**
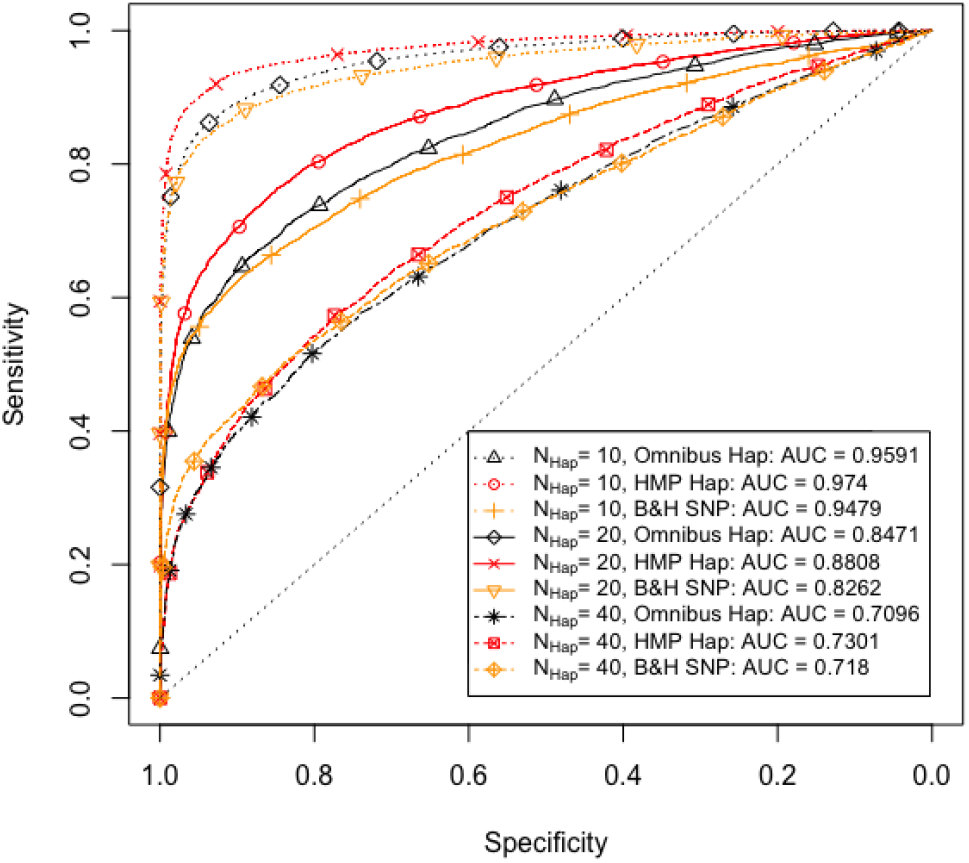
Results for 3 scenarios with different numbers of haplotypes. ROC curves of both haplotype and SNP based tests are shown for 10, 20 and 40 total haplotypes, with all other parameters default as outlined in Table 2.

As detailed in Section 2.4, in scenarios with a large number of haplotypes the starting haplotype frequency is very small and thus the probability that the selected haplotype is lost due to drift in the early phase of the experiment is high. On the other hand, when a large number of haplotypes has high fitness and their fitness values are identical, the chance that the neutral haplotypes are lost in the beginning of the experiment is high. This leads to a scenario where all haplotypes have the same selective strength, and thus there is no selective advantage for any haplotype, resulting in a low power of all considered tests.

We show in Figure S2, that despite a small initial increase, more selected haplotypes will result in lower power for all tests. For the haplotype based tests, the two multiple testing corrections outperform each other in different situations. While HMP performs best with very few and with many selected haplotypes, the omnibus test works best with an intermediate number of selected haplotypes.

Sections S.6.2, S.6.3, S.6.4 and S.6.5 provide simulation results for different values of the number of replicates, the selective strength, number of generations, and the effective population size. As has been previously observed with SNP based tests [43], these results confirm that more replicate populations, a higher selective strength, more generations, and a higher effective population size lead to an increase in power for all tests. The presence of more replicates especially, benefits haplotype based test much more in terms of power compared to the SNP based test. We also observed that the haplotype based tests perform better than the SNP based approach under all considered scenarios.

As our final analysis, we studied the effect of the number of SNPs in the considered window. As shown in Section S.6.6, the haplotype based test is invariant with respect to this parameter. On the other hand, the power of the SNP based test decreases with an increasing number of SNPs. This is due to the more stringent multiple testing correction needed. So in principle, the advantage of haplotype based testing increases with the window size. Notice however, that recombination will become non-negligible with haplotype based testing and will increase the number of relevant haplotypes, making haplotype based testing unfeasible on too large windows.

In experimental evolution, haplotype frequencies are often unknown. If estimates are used instead, and their errors are non-negligible, we propose to use the test statistics introduced in Sections S.2.1 and S.2.2, and S.2.3 instead. They account for the additional variance incurred by haplotype frequency estimates. Both SNP and haplotype based testing will need to take the additional variance into account, if the exact allele frequencies are unknown and replaced by estimates. Notice however, that pool-sequencing noise can be reduced with the haplotype based approach as it ia able to combine the information across SNPs using regression. In Section S.7.2, we consider cases where all frequencies are estimated with a sample size of 500, and with various sequencing coverage values between 50 and 450. With data obtained via pool sequencing, the power decreases only for SNP based testing. Especially for low sequencing coverage, haplotype based testing therefore provides a considerable advantage.

Since both drift and sampling variance affect all tests in similar ways, sampling variation will decrease power. Section S.7.1 provides an illustrative example.

### 3.4 Diploid populations

Here, we evaluate the performance of our proposed methods on a diploid population instead of a haploid one. The population is simulated using the software [49] with 1,000 simulated samples, and other parameters outlined in Table 2. Additive genetic effects are assumed. The results for the diploid population (Figure 3) are similar to the haploid case (Figure 2), where the haplotype based tests outperform the SNP based test in terms of AUC. The haplotype based test using HMP as p-value combination method performs best in line with previously seen haploid results with one selected haplotype. Since heterozygous individuals with one selected allele have a selective advantage of only *s/*2, the power of all tests is lower compared to the haploid case.

**Figure 3:**
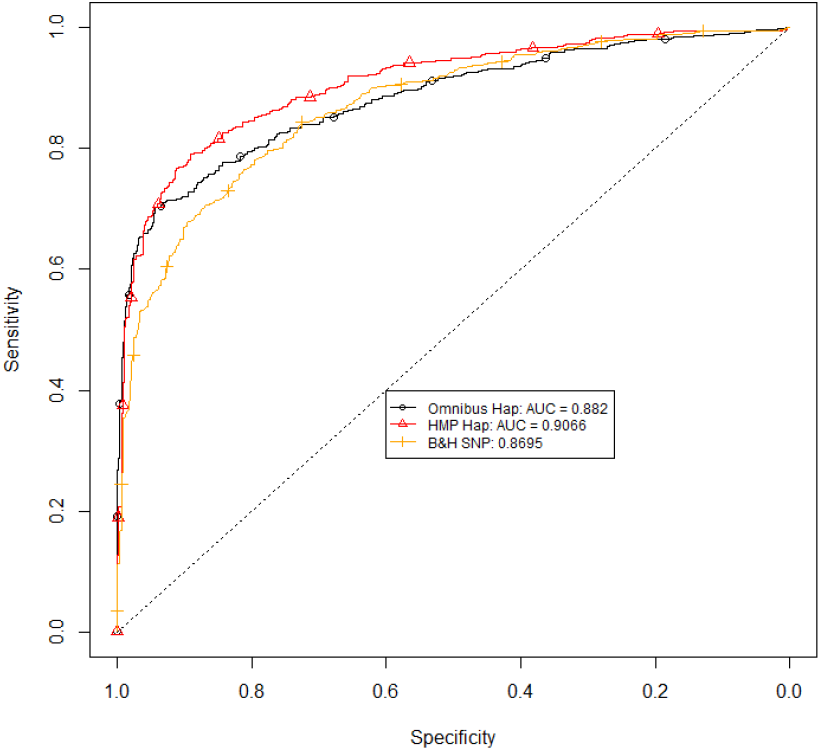
Results for a diploid population. ROC curves of both haplotype and SNP based test are shown for a diploid population. Here, the number of simulations is reduced to 1,000, with all other parameters outlined in Table 2.

### 3.5 Testing for the number of selected haplotypes

The knowledge of the number of selected haplotypes when selection is present is of interest in practical applications when researchers try to better understand the genomic architecture of adaptation in experimental evolution. To investigate the practical performance of the test proposed in Section 2.2, we simulated a scenario with 5 founder haplotypes, some of them selected with *s* = 0.05, that otherwise follows the population parameters in Table 2. We generate 10,000 simulation runs for each considered number of selected haplotypes. If selection is detected by our haplotype based test with the omnibus p-value combination method, we apply our proposed iterative test from Section 2.2.

Figure 4 illustrates that this test is able to accurately predict the number of selected haplotypes for scenarios with different true numbers of selected haplotypes given that selection is strong enough. The left panel presents results conditional on the presence of selection, and the right panel unconditionally.

**Figure 4:**
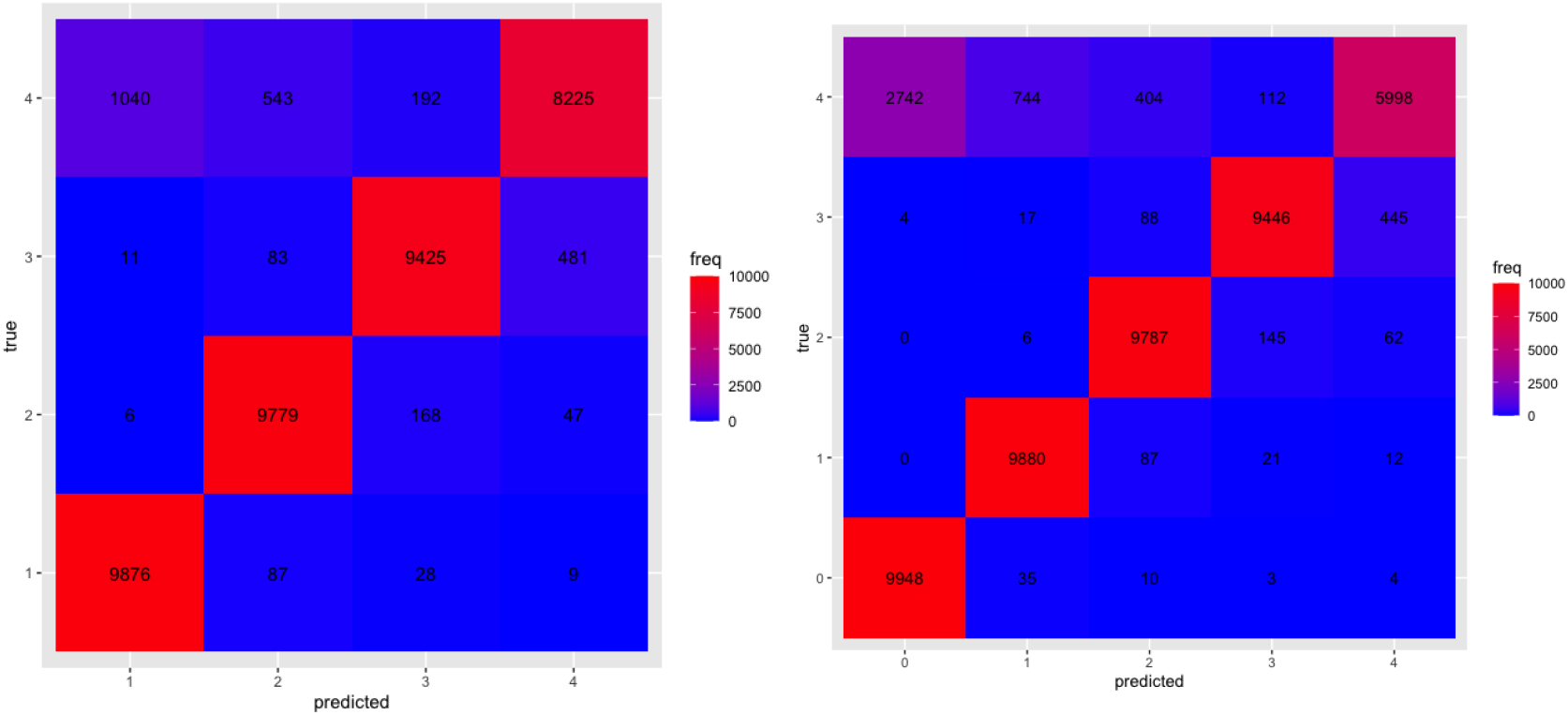
Results showing the prediction accuracy of haplotype based iterative testing. The omnibus method is used for multiple testing correction in all scenarios. The heat map has been obtained using 5 founder haplotypes and a selection strength of *s* = 0.05, with all other parameters default as outlined in Table 2. In the left panel, the haplotype with the strongest signal is always removed as selected. In the right panel, the test is unconditional.

The scenario with one selected haplotype, for instance, is identified correctly in more than 98% of cases both conditionally and unconditionally. We show similar results using the HMP combination method in Section S.9.

### 3.6 Pairwise test

Under the simulation scenarios detailed in Section 3.5, we also explored the performance of the pairwise post hoc test statistic for differences in fitness proposed in Section 2.3.

Figure 5 provides results on the power and the type I error probability, conditional on the rejection of the initial test. Both under the scenarios involving one and three replicate populations, the power of predicting fitness differences between haplotype pairs is high and the type I error probability is controlled. Again, the power is higher with more replicate populations. We observe a decrease in power when the number of selected haplotypes increases.

**Figure 5:**
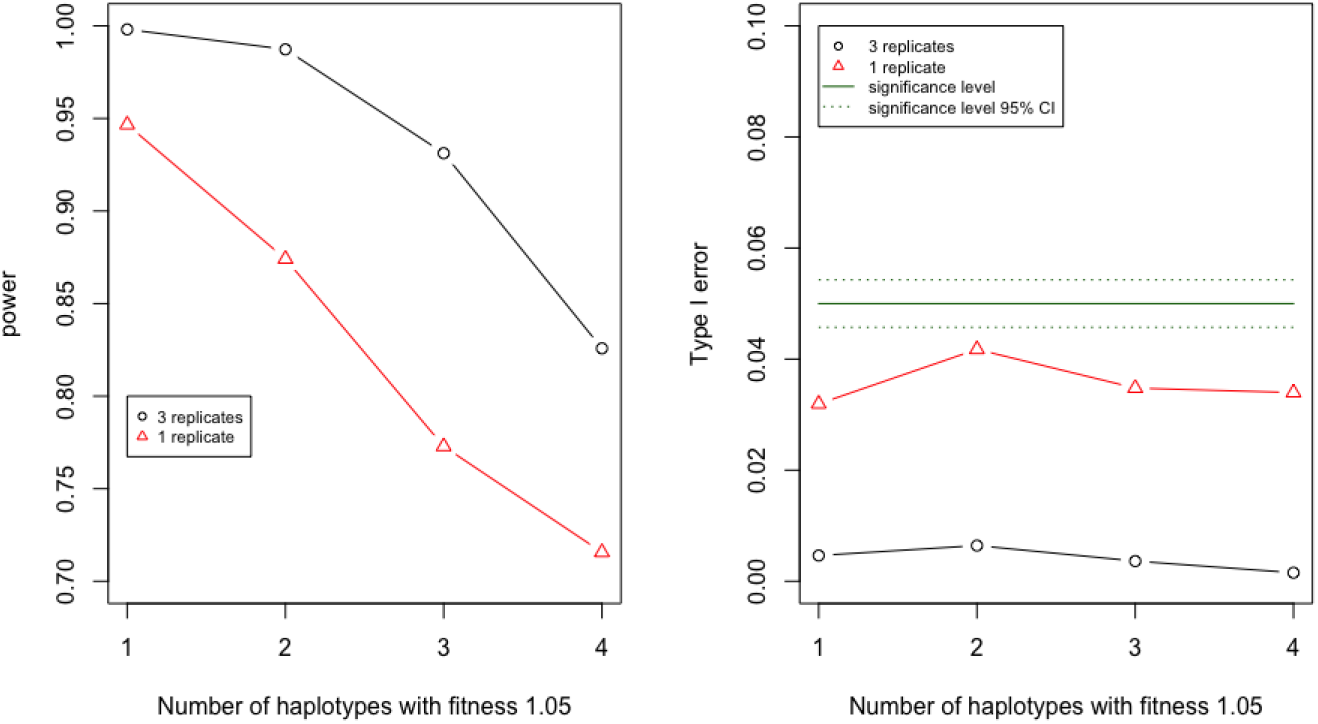
Pairwise tests for differences in fitness for different numbers of selected haplotypes. The displayed probabilities are conditional on the rejection of the initial test for selection. Results based on simulated data using 5 founder haplotypes, selection strength *s* = 0.05, different numbers of selected haplotypes, and 1 or 3 replicate populations, with all other parameters default as outlined in Table 2. Type I errors occur when the test between two haplotypes of equal fitness rejects the null hypothesis. B&H is used for multiple testing correction.

We also investigate the behaviour of the pairwise test under a scenario where all haplotypes differ in fitness. We assign fitness values of 1, 1.02, 1.04, 1.06, and 1.08 respectively to the five considered haplotypes. As one might expect, Figure S18 indicates that power increases with an increasing difference in fitness between the tested haplotypes. For more details on this parameter set, see Section S.11 and S.12.

### 3.7 Experimental designs involving many haplotypes

As discussed already in Section 3.3, the power of our approach decreases with an increasing number of haplotypes. Here we investigate whether the methods proposed in Section 2.4 to reduce the number of haplotypes helps to resolve this problem.

To better understand the interplay between the number of founder haplotypes and that of selected haplotypes, we plot results with an intermediate number of selected haplotypes for different numbers of founders in the left panel of Figure 6. There, *h*_*Sel*_ = *N*_*Hap*_*/*2, while the rest of the parameters are as in Figure 2. As suggested by the results in Figure 6, the haplotype based tests lose their advantage compared to SNP based tests already at 20 founder haplotypes instead of around 40 in Figure 2. Indeed, if many haplotypes have a similar selective advantage, the signal that can be captured by the haplotype based test is diluted. This emphasises that haplotypes should be combined as explained in Section 2.4.

**Figure 6:**
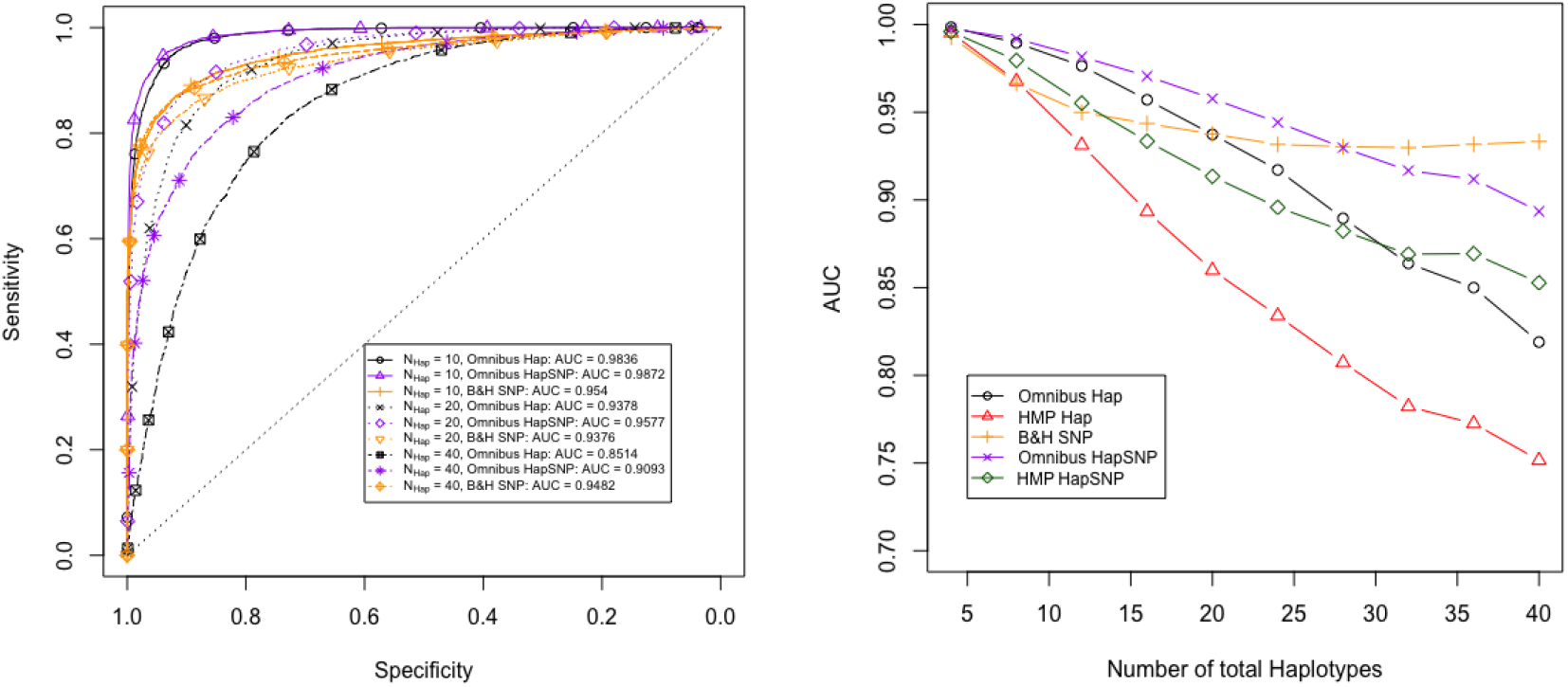
Results for different total numbers of haplotypes. In the left panel, ROC curves are plotted for the original haplotype, HapSNP, and SNP based test under various choices of *N*_*Hap*_. HapSNP refers to the method of using the SNP based test to reduce the haplotype number prior to haplotype based testing outlined in Section 2.4. Here, results are shown for 10, 20, and 40 starting haplotypes. The number of selected haplotypes is set to *h*_*Sel*_ = *N*_*Hap*_*/*2. All other parameter values can be found in Table 2. In the right panel, the AUC of the ROC curves is plotted in dependence of the starting haplotype number. The other parameter values identical to those used in the left panel.

We see that the SNP based combination method (HapSNP) is still able to retain an AUC advantage at 20 haplotypes. Both panels of Figure 6 also show that the SNP based combination method improves the power of the considered haplotype based test consistently. The advantage of haplotype based testing is retained this way also for designs involving a considerably larger number of haplotypes compared to tests that do not operate on a reduced set of haplotypes. The SNP based combination method first reduces the number of haplotypes by removing SNPs, and then applies our proposed haplotype based test.

This approach only requires a minimal increase in terms of computational cost and performs well when the SNP based test is able to identify a reasonable number of individually significant SNPs. When this is not the case, we propose a more computationally demanding modification. This approach reduces the number of haplotypes by creating haplotype blocks and then performs the test for selection on the haplotype blocks. Section S.13 provides results for a scenario with 100 haplotypes, 50 selected haplotypes, and only one population. Here, the haplotype based test combined with the haplotype block based approach performs equally well or better than SNP based tests despite the large haplotype number. This is due to the haplotype block based test being mostly invariant to changes in the number of haplotypes (see Section S.14) Figure S24 provides an example where 30 haplotypes are present, one of them selected. We notice that the SNP based combination method does not perform as well here, in terms of power in subsequent testing. To investigate this further, we considered several significance thresholds which lead to different sets of considered SNPs. However, Figure S25 shows that the choice of threshold seems to have little impact on the performance.

### 3.8 Type I error control

As can be seen in Section S.6, haplotype based tests provide well controlled type I error probabilities, in most of the scenarios we explored. While most changes in parameter value have little to no effect on the type I error of the test, number of replicates, number of haplotypes, and whether the frequencies are known or estimated can have a noticeable impact.

The largest violation in terms of type I error we encountered during our extensive simulations was with the combination of 1 population, 3 to 4 haplotypes, and known haplotype frequencies (Figure S1). Even for these scenarios, the type I error probabilities stay below 0.07, given a significance threshold of 0.05 for the p-values.

We also notice that the CMH test is more conservative than the chi-squared test, causing a drastic decrease in type I error probability when replicates are present. This conservativeness is explored in further detail in Section S.12.

The post hoc tests share similar characteristics in terms of type I error probabilities. The pairwise test tends to have a lower type I error probability compared to the initial test preceding the post hoc procedure (Figure 5), whilst the test for the number of selected haplotypes tends to have a somewhat higher type I error probabilities than the initial test (Figure 4). Indeed, for the pairwise tests we did not observe any violations of the level *α* even under the most liberal scenario. On the other hand, Figure S14 suggests that the true number of haplotypes is over-estimated in about 11% of cases at worst.

One scenario when such a claim does not necessarily hold is due to the type I error being computed conditional on the rejection of the null hypothesis of the haplotype based test. These conditional type I errors often have higher probabilities compared to unconditional type I errors, which are not dependent on the rejection of the null hypothesis. This is the case in particular under weak selection, where a larger proportion of the initial rejections is caused by inflated effect sizes due to random errors. Due to such a filtering, the conditional type I error can exceed the desired threshold, while the unconditional type I error is well under control. With one selected haplotype, Figure S17 provides an example that shows that the conditional type I error is not under control under a very low selective strength *s* = 0.01.

With the haplotype block based tests there are two levels of multiple testing correction, one within and one between the haplotype blocks. We suggest the usage of HMP or B&H as multiple testing correction at the second (between) layer, since similarities between the blocks can introduce a fairly strong positive correlation among the tests.

## 4 Real data application

Here we illustrate our proposed testing approach on yeast data from [35], where both haplotype and allele frequencies have been obtained at 3 different time points. The experiment involves 4 haplotypes which are investigated on a grid of non-overlapping 30KB windows. We tried the omnibus variant of the haplotype based test, as well as SNP based testing using a within-window Benjamini & Hochberg correction. For both methods, we also applied a Benjamini & Hochberg multiple testing correction across windows in order to control a false discovery rate of 0.05 at a genome-wide level. The three sequenced time points have been taken at cycles 0, 6 and 12, where each cycle is estimated to contain 15 to 20 generations. For our analysis, we take the mean and assume 17.5 generations per cycle which leads to 105 generations between adjacent sequenced time points.

We estimated the effective population size with the method proposed by [36]. Here, we use data from population 4k, at cycles 0 and 6. We did not include the frequencies at cycle 12 due to the high percentage of fixed or lost alleles, and the very low estimated effective population sizes, see Section S.16 for more details. We also check windows with very highly correlated haplotypes by constructing UPGMA trees. Such windows may lead to false positive results due to multi-collinearity, when assuming that the error in haplotype reconstruction is negligible. We did not find this to be an issue for the windows we detect as selected. Table 3 provides the percentage of windows each method detects as affected by selection separately per chromosome. The absolute number of windows detected can be found in Section S.17.

**Table 3:** Percentage of windows each method is able to detect as selected for chromosomes 1-16.

| Chromosomes | Total | 1 | 2 | 3 | 4 | 5 | 6 | 7 | 8 | 9 | 10 | 11 | 12 | 13 | 14 | 15 | 16 |
| --- | --- | --- | --- | --- | --- | --- | --- | --- | --- | --- | --- | --- | --- | --- | --- | --- | --- |
| Hap based | 0.133 | 0.129 | 0.049 | 0.211 | 0.140 | 0.331 | 0.096 | 0.040 | 0.028 | 0.098 | 0.281 | 0.215 | 0.077 | 0.110 | 0.172 | 0.111 | 0.149 |
| SNP based | 0.070 | 0.188 | 0.022 | 0.289 | 0.098 | 0.000 | 0.000 | 0.150 | 0.176 | 0.000 | 0.058 | 0.000 | 0.049 | 0.060 | 0.011 | 0.049 | 0.070 |

For 12 out of the 16 chromosomes and also genome-wide, the haplotype based test detects more selected windows than the SNP based test. This is in line with our previous simulations that confirmed a higher power of haplotype based tests under most considered scenarios.

To better understand the selective architecture, we next investigate the number of selected haplotypes for the 39 windows on chromosome 9 which we identified as affected by selection. Out of the 39 windows, one selected haplotype was identified for 3 windows. Furthermore, 35 (resp. 1) windows led to a test result suggesting 2 (resp. 3) selected haplotypes.

Figure 7 displays a genome-wide summary of the selected window positions and the respectively inferred numbers of selected haplotypes using our proposed haplotype based testing approach. On chromosome 6, for instance, more than 1 selected haplotype is identified for all selected windows.

**Figure 7:**
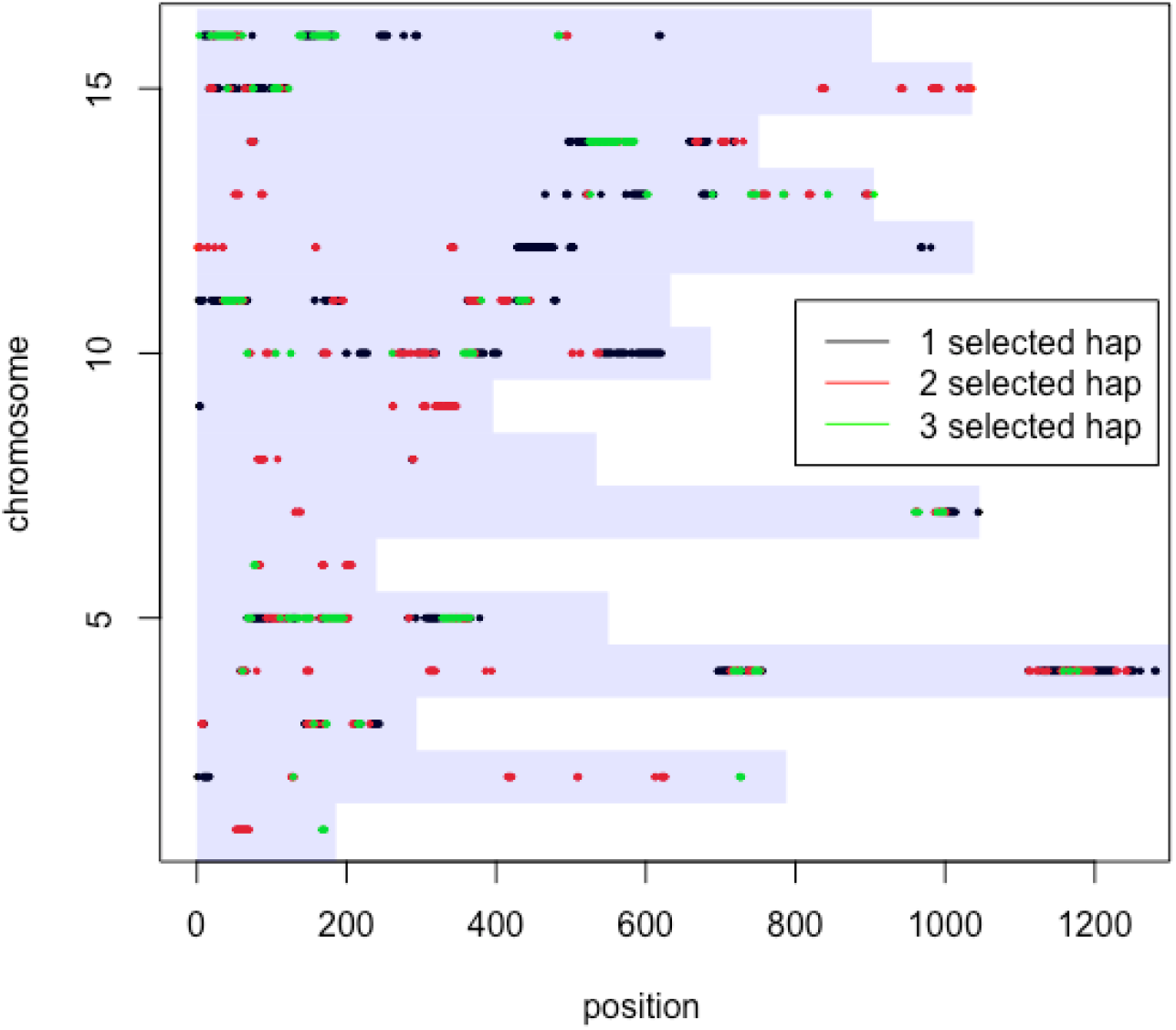
Position of selected windows according to haplotype based test. The background rectangles represent the chromosomes, the positions represent the window positions and can be translated into positions on the genome when multiplying them by 30KB. The dots represent positions of selected windows found by the (omnibus) haplotype based test, and the colours indicate the number of selected haplotypes inferred by our proposed post hoc test.

We also see that some selected windows cluster together which could point towards hitchhiking effects in neighbouring windows. Similar numbers of selected haplotypes for nearby windows, may also point towards selected haplotypes extending over multiple windows.

We also applied pairwise tests to chromosomes 9 and 6, and summarise the results in Table 4, with haplotypes H1, H2, H3, H4 denoting the haplotypes alt YEE hap A1 00, alt YEE hap A2 00, alt YEE hap B3 00, alt YEE hap B4 00 respectively.

**Table 4:** Proportion of windows for which the pairwise tests provide evidence of fitness differences between specific haplotype pairs.

| Haplotype pairs<br>Chromosome | H1-H2 | H1-H3 | H1-H4 | H2-H3 | H2-H4 | H3-H4 |
| --- | --- | --- | --- | --- | --- | --- |
| 6 | 0.130 | 1.000 | 0.000 | 0.783 | 0.000 | 1.000 |
| 9 | 0.154 | 0.821 | 0.000 | 0.692 | 0.154 | 0.846 |

A further inspection of the outcome for chromosome 6 suggests that H1 and H4 are positively selected for all selected windows according to our test. H3 seems to be the lowest fitness haplotype, and H2 shows lower fitness compared to H3 for some selected windows.

### 4.1 Simulation study using parameters from real dataset

To further validate our results from our real data example, we carried out simulations using the haplotype structure matrix, the starting frequencies, sample size, estimated *N*_*e*_, number of generations and number of selected haplotypes from this example.

We looked at the windows where we were able to detect selection using the haplotype based test, but not with the SNP based one.

Figure 8 focuses on chromosome 9 window 261, where two selected haplotypes were found with our method. The window consists of 457 SNPs, an average sample size of 92, and is estimated to have a *N*_*e*_ of 1916.

**Figure 8:**
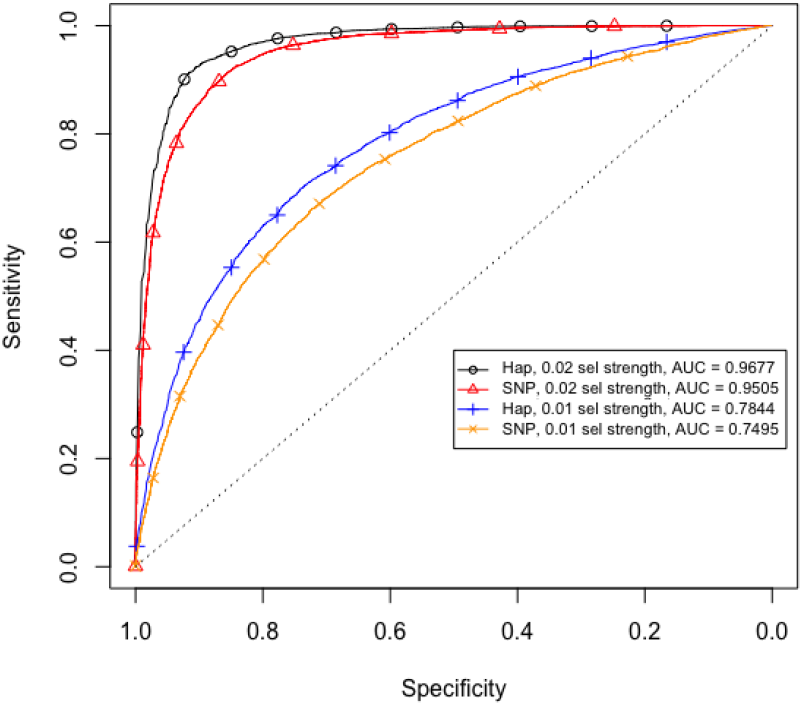
Results based on a simulation using parameters of chromosome 9, window 261. ROC curves of both haplotype and SNP based test are shown under selection strengths 0.01 and 0.02.

We see that according to the simulation, the haplotype based test performs consistently better in terms of AUC at both selective strengths, which might explain why we were able to detect selection at this window only with the haplotype based test. Type I error is also controlled for both haplotype and SNP based tests under given parameter settings.

## 5 Conclusion

It is well known that haplotypes harbour valuable information that can be helpful for the understanding of the selective architecture [45]. In this work, we show that haplotype based tests can effectively detect selection under a wide range of evolve and resequence scenarios. In order to apply haplotype based tests, we focus on relatively short genomic windows such that the haplotypes are not affected by recombination and their frequency changes can be followed over time. In [17] the window size for various organisms is provided such that the impact of recombination can be assumed negligible, which may be used as a guideline. We demonstrate that haplotype based tests are more powerful than traditional SNP based approaches for E&R experiments when the number of haplotypes is small to moderate. Since such genomic regions can include several hundreds of SNPs and usually much fewer haplotypes, the required multiple testing correction is much more severe at a SNP level than for haplotype based tests.

When selection has been detected, we also propose post hoc tests to infer the number of selected haplotypes, as well as haplotype pairs that differ in fitness. These post hoc tests may contribute to a better understanding of the selective architecture. They perform especially well when data from replicate populations showing consistent signals are available.

For experimental designs involving too many haplotypes for efficient testing, we propose methods of collapsing their frequencies. These methods either use SNPs identified by preliminary tests, or a haplotype block based approach. They help to reduce the loss in power incurred with an increasing number of haplotypes and increase the range of scenarios for which haplotype based tests are a good choice. In experiments involving large numbers of haplotypes (say above 100), approaches where the results based on haplotypes and on SNPs are combined would be an interesting follow up.

We studied our proposed haplotype based test under a wide range of simulated experiments. The variables that mostly influence the performance of the test are the number of haplotypes, and selected haplotypes. These parameters also affect the power of the SNP based tests. The size of the considered window is also relevant for SNP based tests, as it influences the amount of required multiple testing corrections. However, it does not affect the power of the haplotype based test, as long as recombination remains negligible. Our investigation of the effect of different design parameters should be useful when planning new experiments, as their choices have an important influence on the power of subsequent tests for selection.

In this paper, most of our simulation results were obtained for scenarios with known SNP and haplotype frequencies. However, often the haplotype frequencies are unknown in applications [20, 40]. Thus we also provide versions of our tests for scenarios with unknown or partially unknown haplotype frequencies.

With sampling errors, the relative performance between haplotype and SNP based tests remains similar to the known frequency scenario. The advantage of haplotype based tests can increase considerably, however, if additional noise is introduced by pool sequencing. The smaller the sequencing coverage, the more noise is added by pool sequencing. Since haplotype frequencies can be estimated by combining information across many SNPs using for instance regression, haplotype based methods are much less affected by pool sequencing noise than approaches focusing on individual SNPs.

Our results demonstrate that haplotype based tests for selection provide attractive tools to better understand selective architectures in the context of experimental evolution. Both the initial and the post hoc tests contribute to such a goal.

We implemented our proposed approach in an R [51] package available at https://github.com/xthchen/haplotest.

## 6 Acknowledgements

Haoyu Chen, Marta Pelizzola and Andreas Futschik acknowledge support of the Austrian Science Fund (FWF; DK W1225-B20). This research was also supported in part by the National Science Foundation under Grant No. NSF PHY-1748958.

## Supplementary material

### S.1 Multiple testing corrections

We have considered using other multiple testing corrections for our haplotype based approach, such as the commonly used Bonferroni [4] and Benjamini & Hochberg [6], as well as Vovk’s methods proposed in [27]. During our preliminary work however, these tests have shown to be much more conservative than omnibus and HMP due to stricter constraints.

#### S.1.1 Vovk’s methods

[27] proposed combination methods that promise type I error control under any dependencies given that the number of p-values is large. The methods are extremely conservative similar to the Bonferroni correction, which is a special case of the proposed methods, but can be useful if having very few false positives is the goal. The methods depend on a parameter value *r*, with guidelines suggesting that more dependent p-values should use larger values of *r*. The merging function dependent on the total number of p-values *N* and parameter *r* is defined as follows:

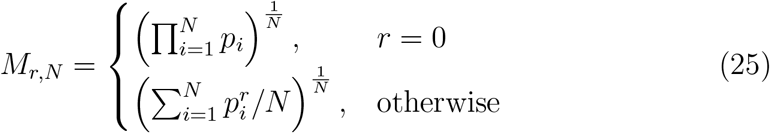

with evidence of rejecting the null hypothesis present if:

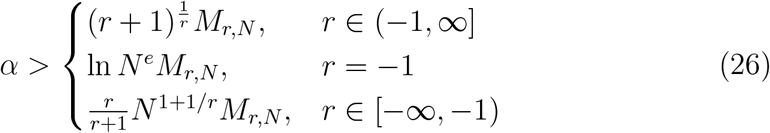

The Bonferroni correction would correspond to the case when *r* = *−∞*.

### S.2 Test statistic under different scenarios

In the paper we have shown the test statistic for haplotype based testing in a scenario where haplotype frequencies are known, and effective population size does not change over time. This however, might not be realistic. Usually, haplotype and allele frequencies are estimated with either or both sampling and pool sequencing noises. When such is the case, variances from these noises need to be taken into account to avoid type I error violations. Here, we provide test statistics for a wider range of scenarios, where the frequencies are estimated with different sources of variance.

#### S.2.1 Test statistic when all haplotype frequencies are estimated with sampling variance

Assuming a sample size of size 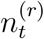 is used to estimate the haplotype frequencies at time *t* and replicate *r*, an additional variance component of frequency estimation is present when compared to the case with known frequencies.

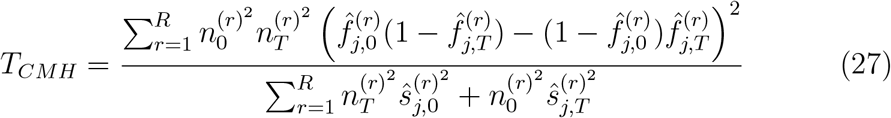

with 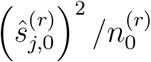 and 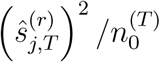 being consistent estimators of 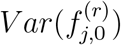 and 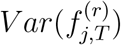 respectively:

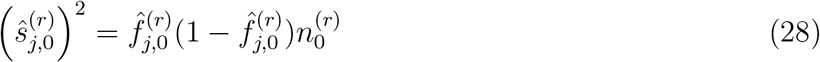

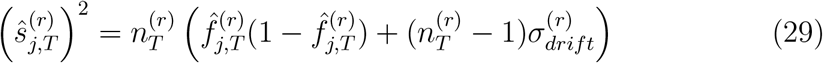

#### S.2.2 Test statistic when starting haplotype frequencies are known, others estimated with sampling variance

Similar to the scenario in Section S.2.1, except the variance component for starting frequencies is nullified.

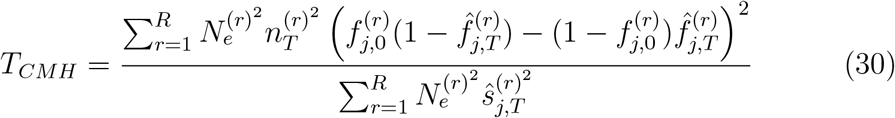

with 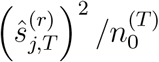 being consistent estimators of 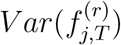:

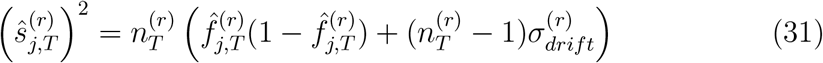

#### S.2.3 Test statistic when all frequencies are estimated with both sampling and pool-sequencing variances

Assuming a sample size of 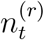 and coverage of 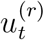 at time *t* and replicate *r*, additional components of variance are needed to be taken into account. The new test statistic under this scenario is given as follows:

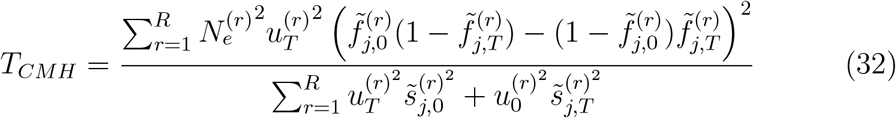

with 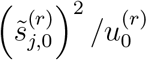 and 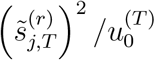 being consistent estimators of 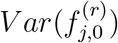 and 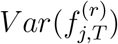 respectively. For SNP based test, this can be computed by:

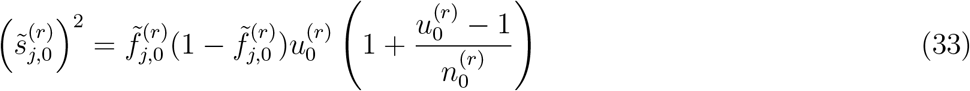

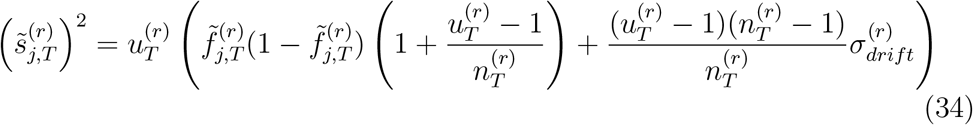

and for haplotype based test, these are the variances of the estimated regression coefficients 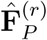 at time points 0 and *T*. Further derivations for the SNP based tests can be found in [7].

#### S.2.4 Drift variance when *N*_*e*_ changes over time

If *N*_*e*_ changes due to experimental design, and is measured at all sequenced timepoints, or due to testing for number of selected haplotypes, the variance of drift with changing *N*_*e*_, 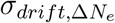 can be computed by:

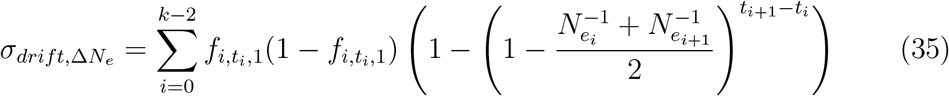

where 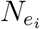 is the *N*_*e*_ at timepoint *t*_*i*_. 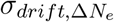 can then be used in place of *σ*_*drift*_ in the previous test statistics.

#### S.2.5 Testing for the number of selected haplotypes when frequencies are estimated with sampling variance

Assuming the presence of *R* replicates and estimated frequencies, the *m*_1_ haplo-type that provides the maximum change in frequency outcome is calculated as follows:

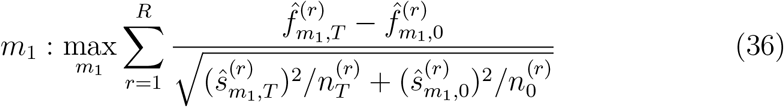

Notice that due to the noise introduced by sequencing and sampling, estimated intermediate frequencies can be zero, while latter frequencies deviate from zero again.

### S.3 Combining haplotypes using SNP based test

As introduced in Section 2.4, by removing certain SNPs, it could be possible that several haplotypes now share the same set of nucleotides, and their frequencies can thus be combined. Even though due to hitchhiking, smaller p-values do not necessarily indicate a selected target, selected targets will unlikely result in large p-values. Thus through the removal of SNPs that give p-values higher than some certain threshold *β*, the combined haplotypes are likely to either be both targets or non-targets of selection, achieving the goal of haplotype number reduction while not diluting the original signal. Below we explain this idea in more detail.

Step 1: From allele frequency matrix 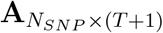, apply adapted chi-square test [7] as in haplotype based test.

Step 2: Using the obtained p-values 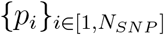, create the filtered set of SNPs 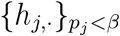.

Step 3: Create the new haplotype structure matrix **H**^*filt*^ by removing all SNPs from **H** not present in the filtered SNP set.

Step 4: Combine all haplotype frequencies that now have the same nucleotide 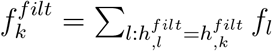.

Step 5: Apply the haplotype based test on the new combined haplotype frequency matrix **F**^*filt*^.

### S.4 Combining haplotypes using haplotype blocks

In this section we provide more details about the computation of haplotype blocks. Suppose in a simple setup where we have 2 SNPs and hence 4 haplotypes, with the allele frequencies of the two SNPs being *u, v* respectively, we write the haplotype frequencies as *h*_1,1_, *h*_1,0_, *h*_0,1_, *h*_0,0_ dependent on the nucleotide they hold. The coefficient of linkage disequilibrium, D, is then calculated by:

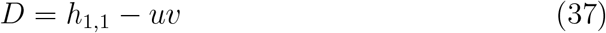

This value is not informative as its size is relative to the allele frequencies, therefore usually a normalised version *D*′ proposed by Lewontin (1964) [29] is used to measure LD:

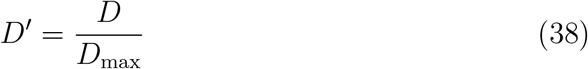

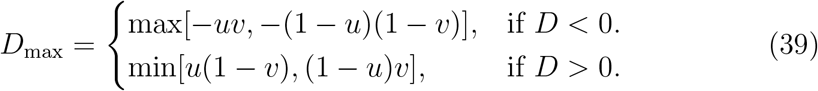

The variance of *D*′ can be estimated as suggested in [30]:

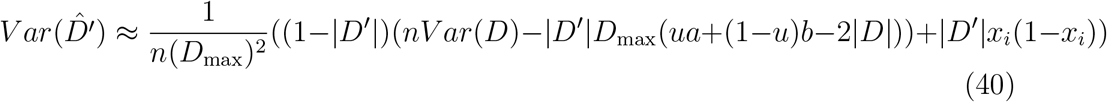

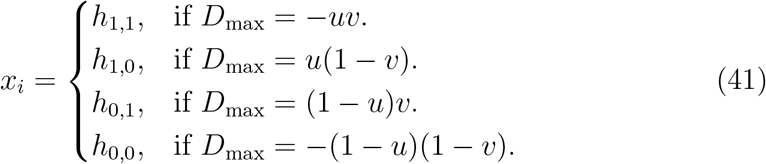

where *n* is the total number of haplotypes of the entire interested SNP region, and *a* = *v, b* = (1 *− v*) if *D*′ *<* 0 and vice versa. If we write the normalised coefficient of linkage disequilibrium between some SNP pairs *i* and *j* as 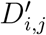, by using this variance approximation and assuming 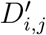 to be normally distributed, we can calculate its 90% confidence interval, with the upper and lower intervals being *U*_*i,j*_ and *L*_*i,j*_ respectively. The definition of an haplotype block is then given by Gabriel et al. (2002) [28], where the pair of SNPs are said to be in strong linkage disequilibrium (LD) if *L*_*i,j*_ *≥* 0.7 and *U*_*i,j*_ *≥* 0.98 and the pair shows strong evidence of historical recombination, or EHR if *U*_*i,j*_ *<* 0.9.

A haplotype block from SNP *i* to SNP *j* can only exist if the SNP pairs *i* and *j* are in strong LD. The number of SNP pairs that are in strong LD within the block needs to be at least 19 times the number of SNP pairs that are in EHR. After finding all possible haplotype block within the given window, we rank the haplotype blocks according to SNP length, and select non-overlapping blocks as our block choice starting from the highest rank. We only use haplotype blocks that extend over more than 2 SNPs, as little information is gained from very short blocks.

### S.5 Type I error control without replicates

**Figure S1:**
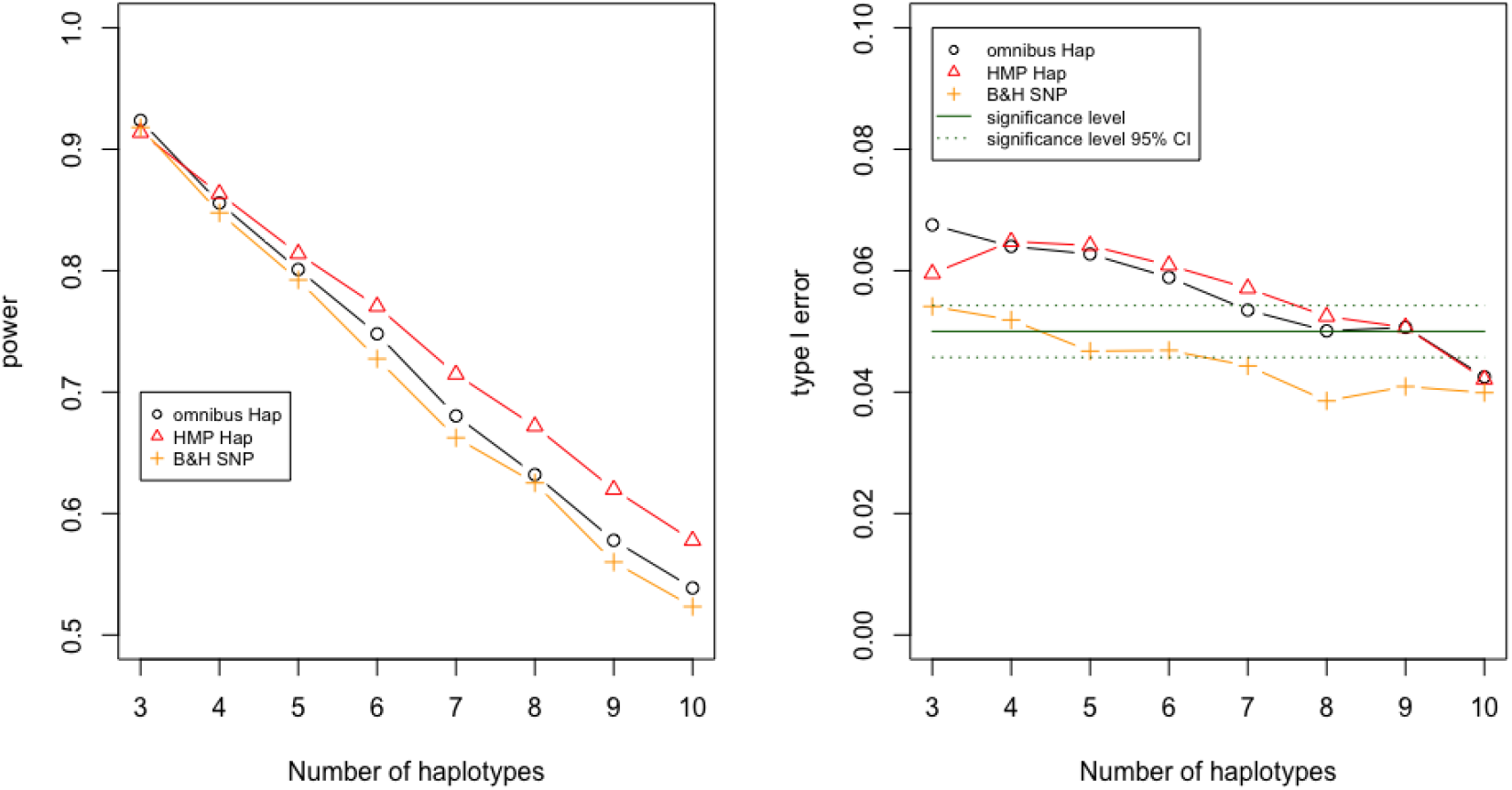
Results when changing the number of total haplotypes with no replicate population. Power and Type I error curves are plotted for both haplotype and SNP based test under various *N*_*Hap*_, no replicate population, with all other parameters default as outlined in Table 2.

### S.6 Effect on power and AUC when changing model parameters

#### S.6.1 Number of selected haplotypes

**Figure S2:**
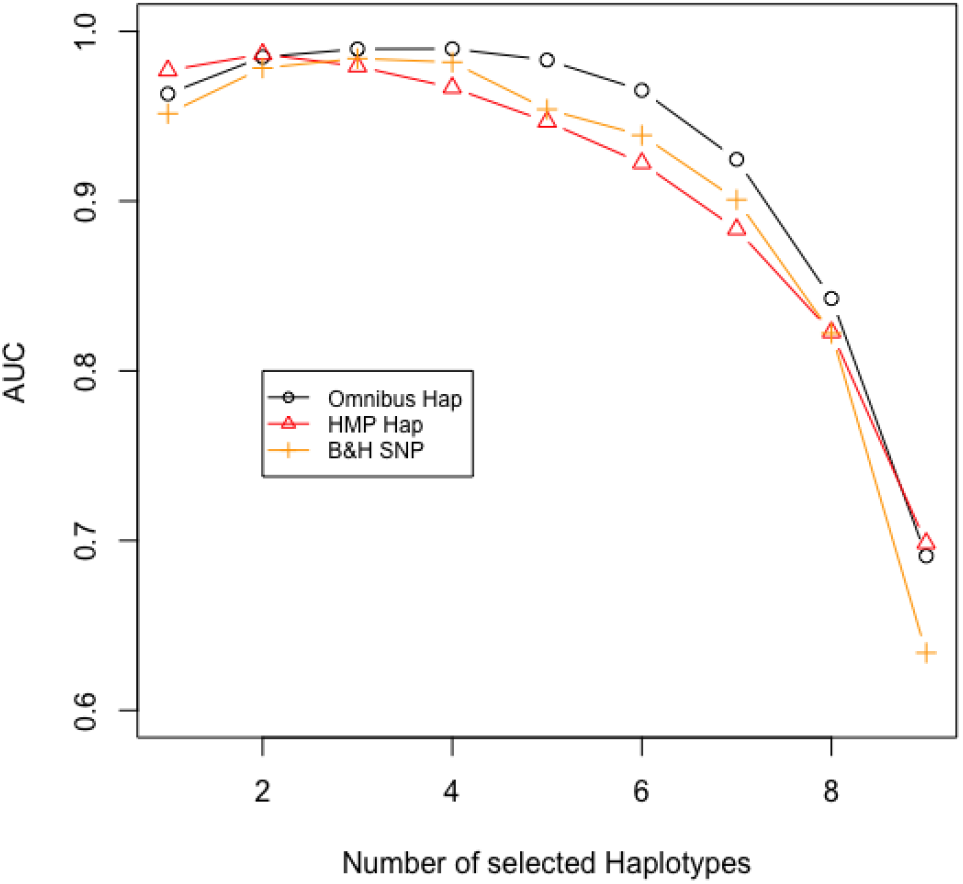
Results when changing the number of selected haplotypes *h*_*Sel*_. Area under curve (AUC) of the ROC curves are provided for the haplotype and SNP based tests under various values of *h*_*Sel*_, with all other parameters default as outlined in Table 2.

#### S.6.2 Number of replicates

**Figure S3:**
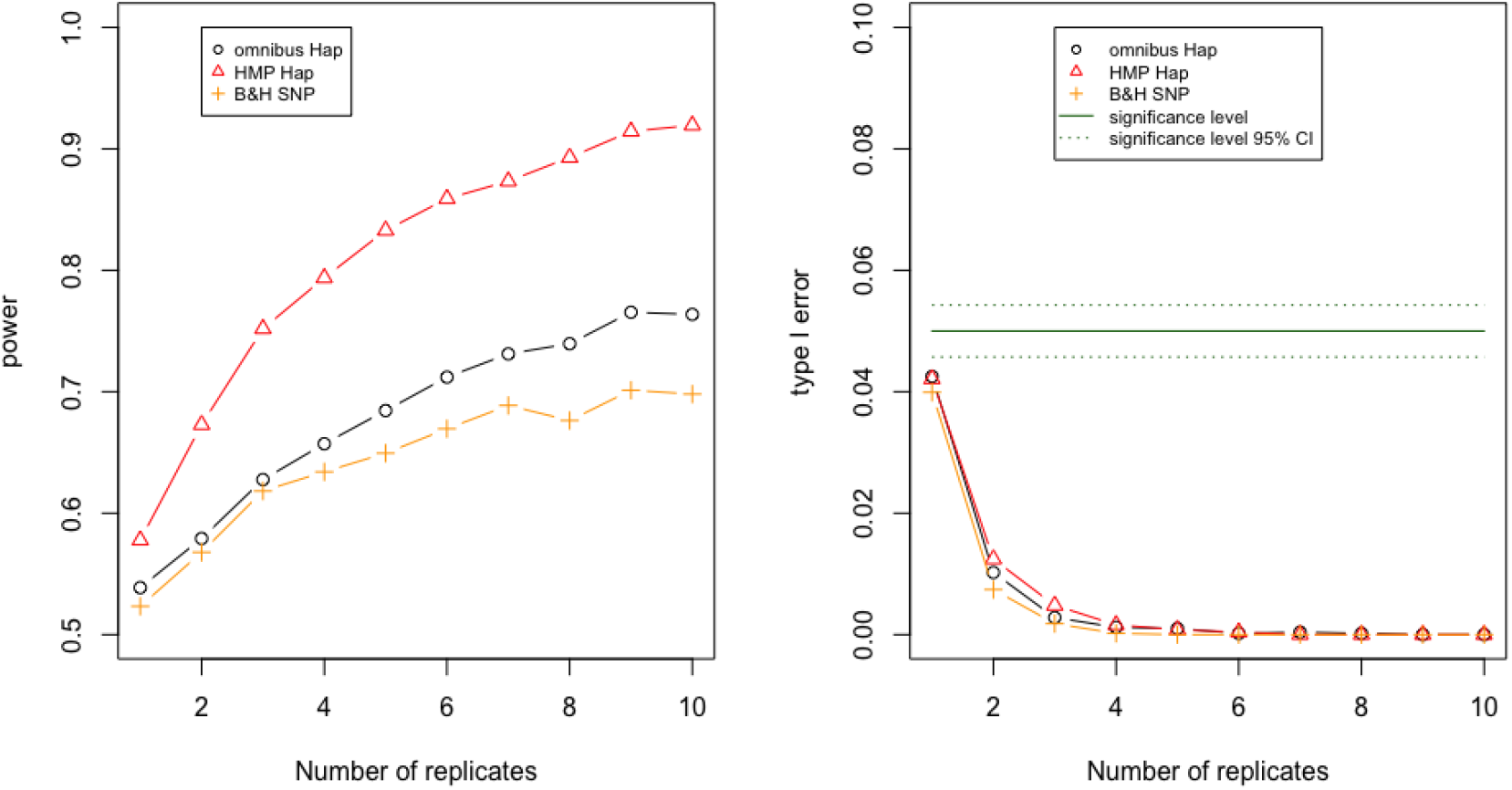
Results when changing the number of replicate populations. Power and Type I error curves are plotted for both haplotype and SNP based test under various values for the number of replicates *R*, with all other parameters default as outlined in Table 2.

#### S.6.3 Selective strength

**Figure S4:**
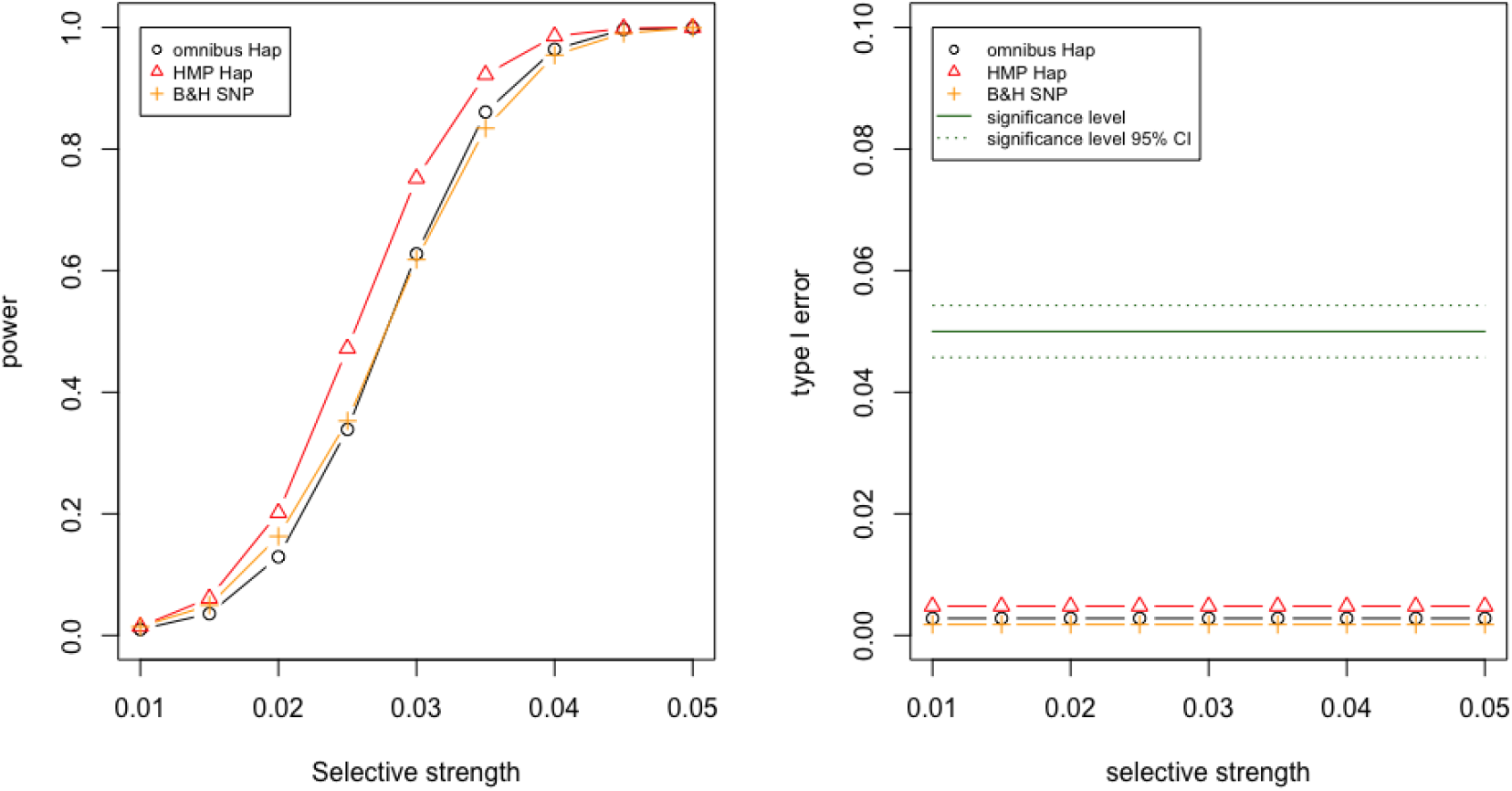
Results when changing selection strength. Power and Type I error curves are plotted for both haplotype and SNP based test under various values for the selection coefficient *s*, with all other parameters default as outlined in Table 2.

**Figure S5:**
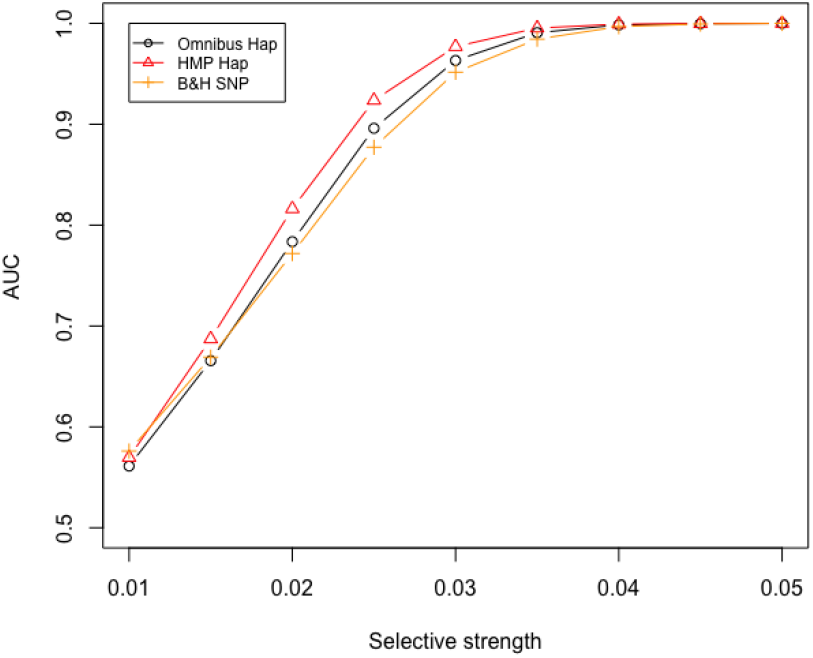
Results when changing selection strength. AUC of the ROC curves are plotted for both haplotype and SNP based test under various values for the selection coefficient *s*, with all other parameters default as outlined in Table 2.

#### S.6.4 Number of generations

**Figure S6:**
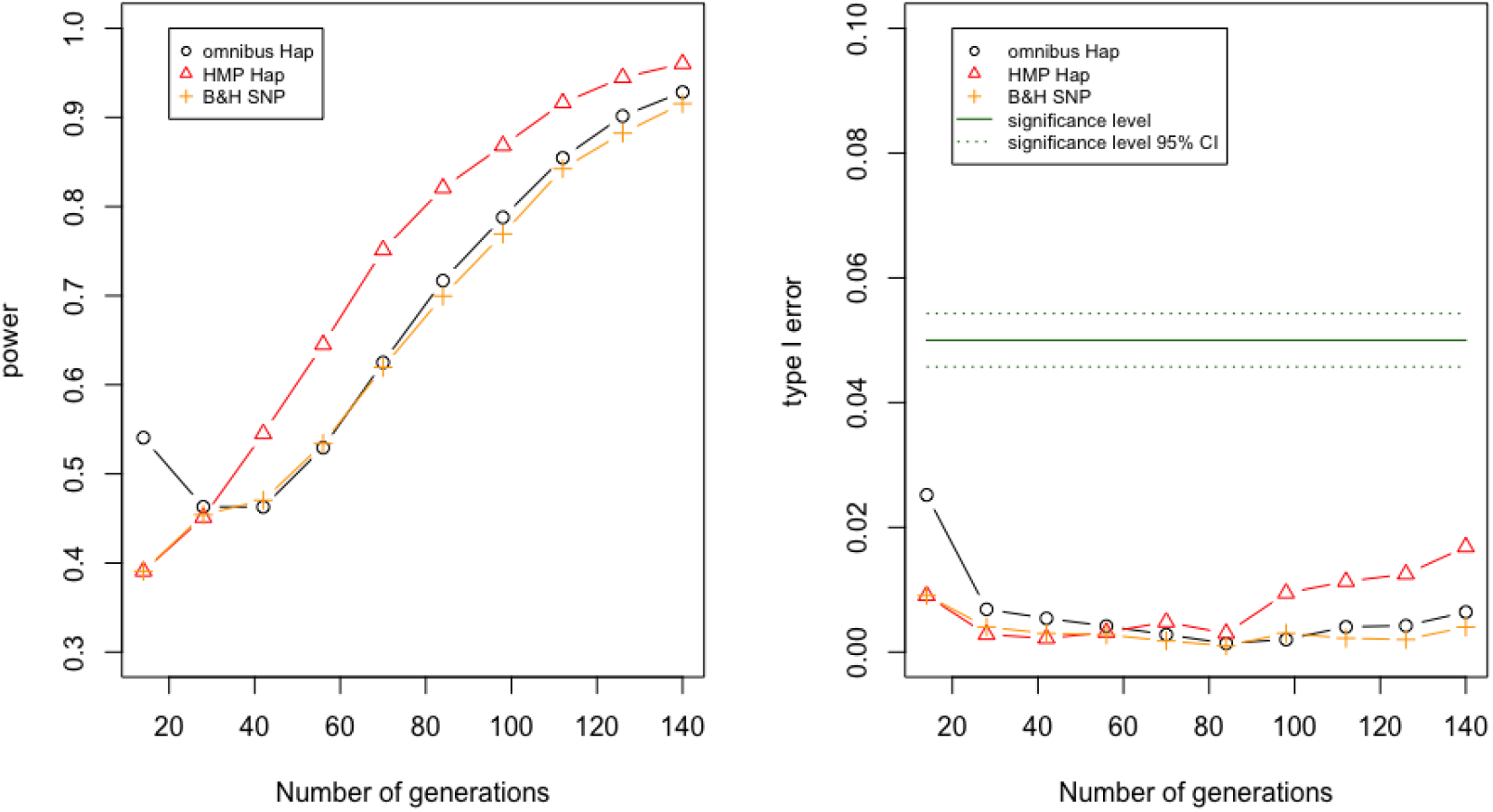
Results when changing the total number of generations. Power and Type I error curves are plotted for both haplotype and SNP based test under various total number of generations, with all other parameters default as outlined in Table 2.

**Figure S7:**
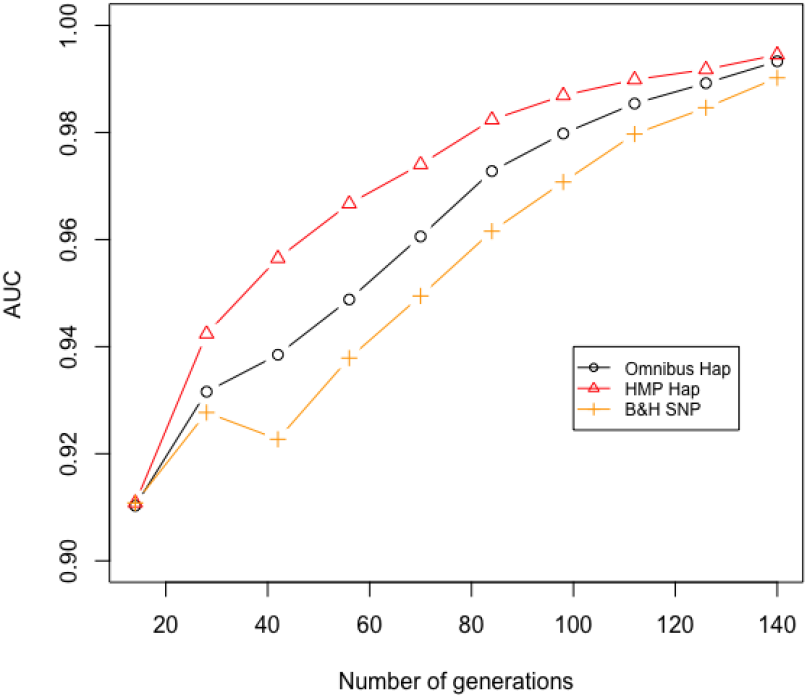
Results when changing the total number of generations. AUC of the ROC curves are plotted for both haplotype and SNP based test under various total number of generations, with all other parameters default as outlined in Table 2.

#### S.6.5 Effective population size

**Figure S8:**
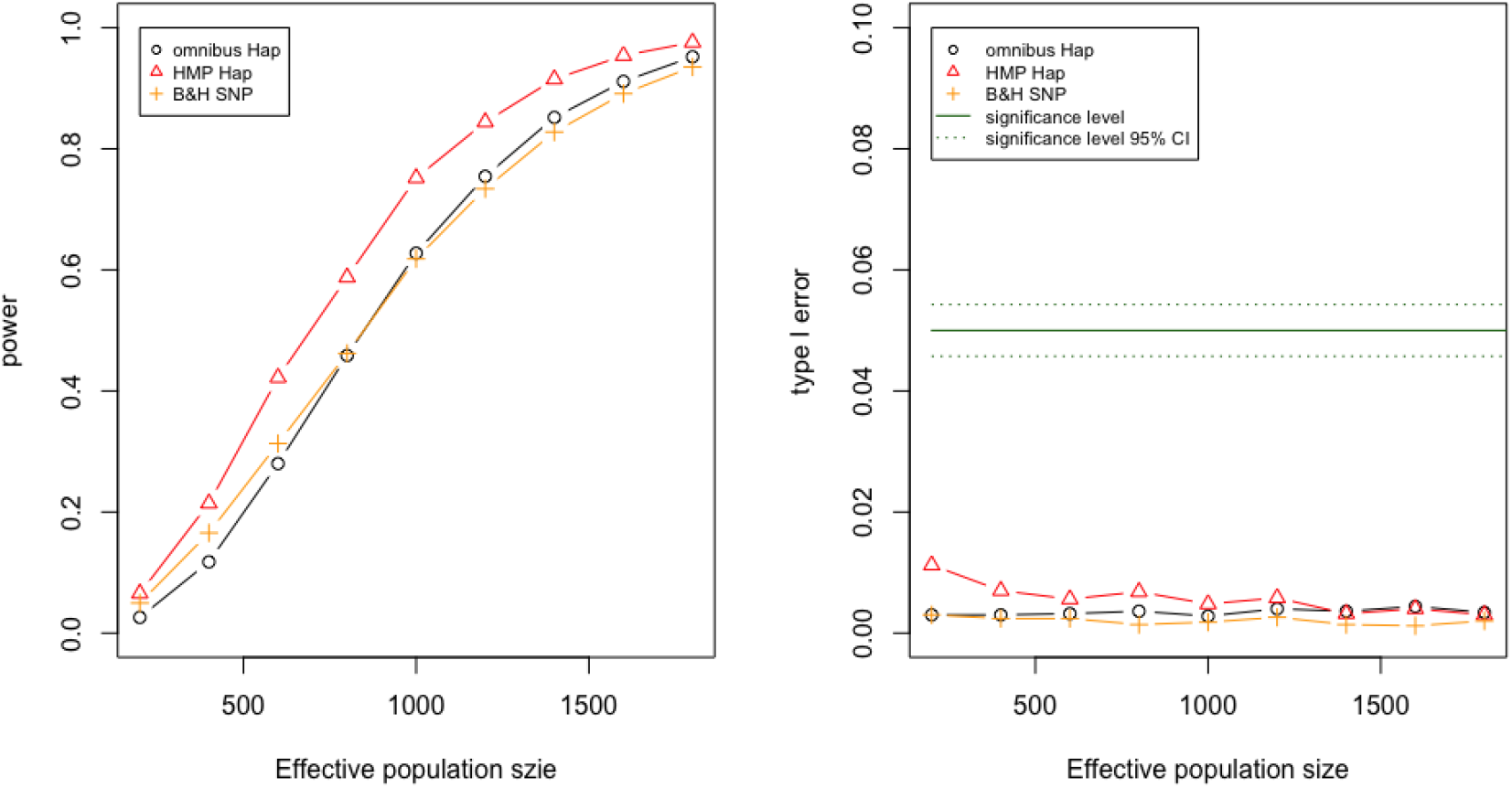
Results when changing effective population size. Power and Type I error curves are plotted for both haplotype and SNP based test under various *N*_*e*_, with all other parameters default as outlined in Table 2.

**Figure S9:**
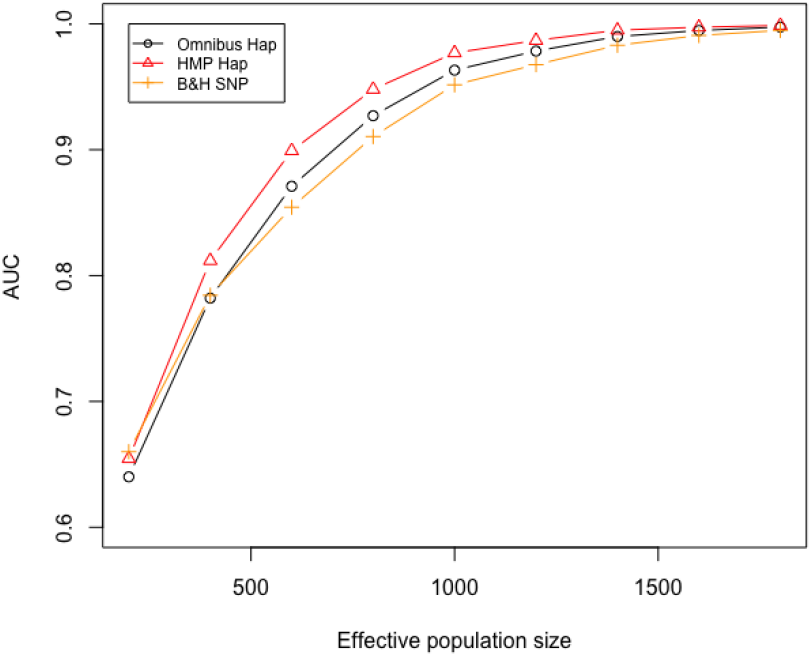
Results when changing effective population size. AUC of the ROC curves are plotted for both haplotype and SNP based test under various *N*_*e*_, with all other parameters default as outlined in Table 2.

#### S.6.6 Number of SNPs

**Figure S10:**
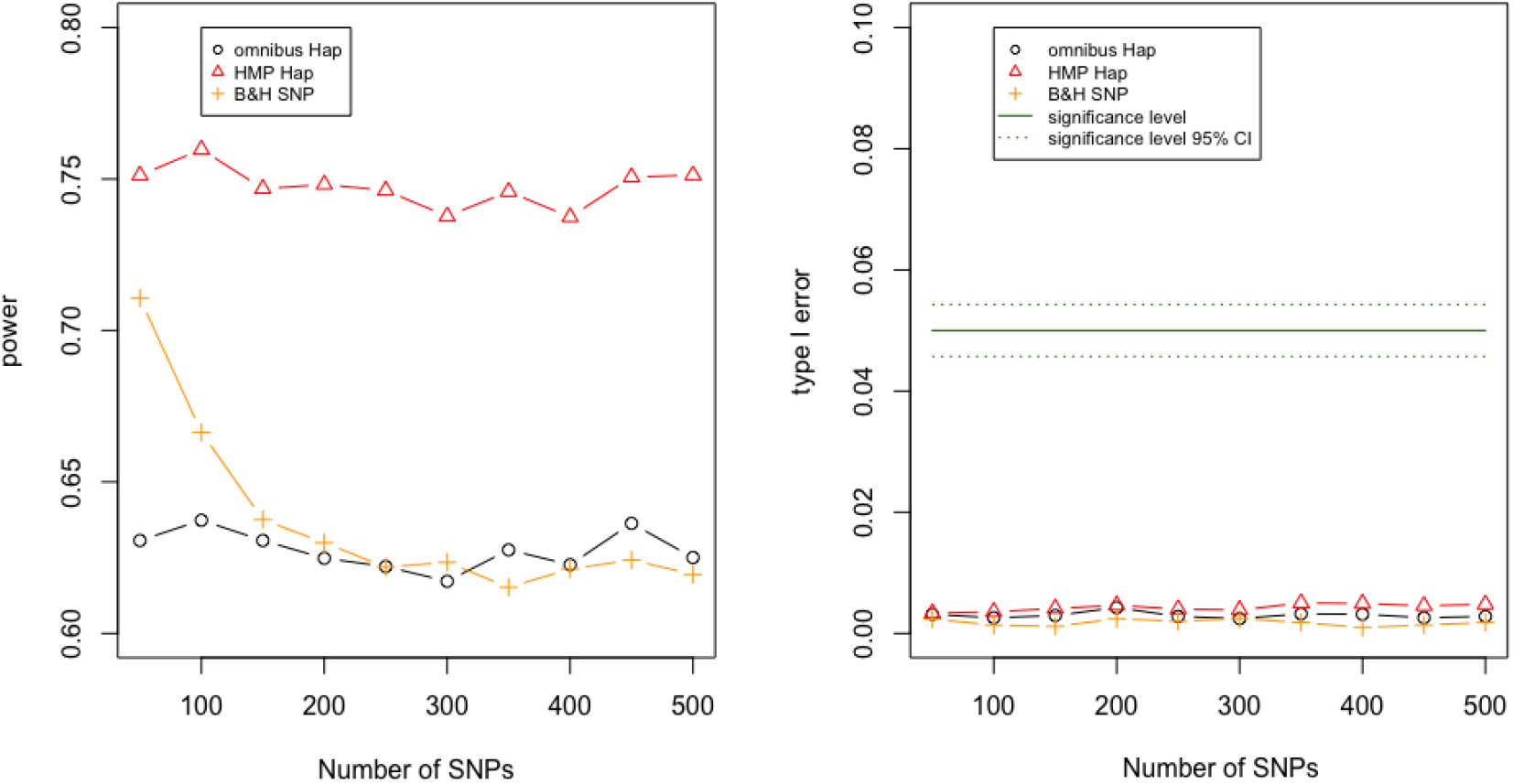
Results when changing the number of SNPs. Power and Type I error curves are plotted for both haplotype and SNP based test under various *N*_*SNP*_, with all other parameters default as outlined in Table 2.

**Figure S11:**
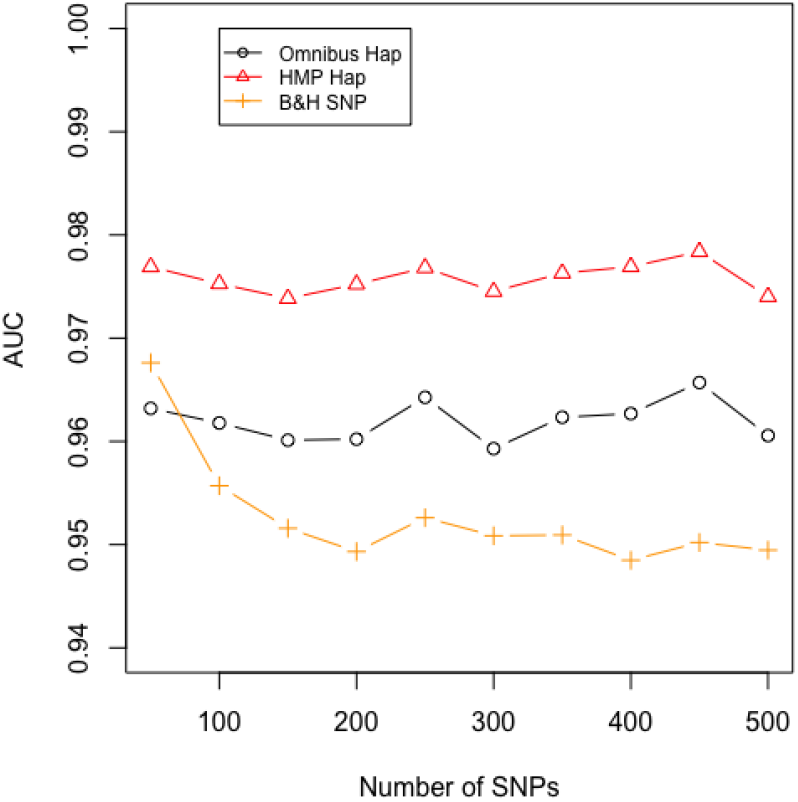
Results when changing the number of SNPs. AUC of the ROC curves are plotted for both haplotype and SNP based test under various *N*_*SNP*_, with all other parameters default as outlined in Table 2.

### S.7 Different scenarios of frequency estimation

#### S.7.1 Power and type I error when sampling noises exists

**Figure S12:**
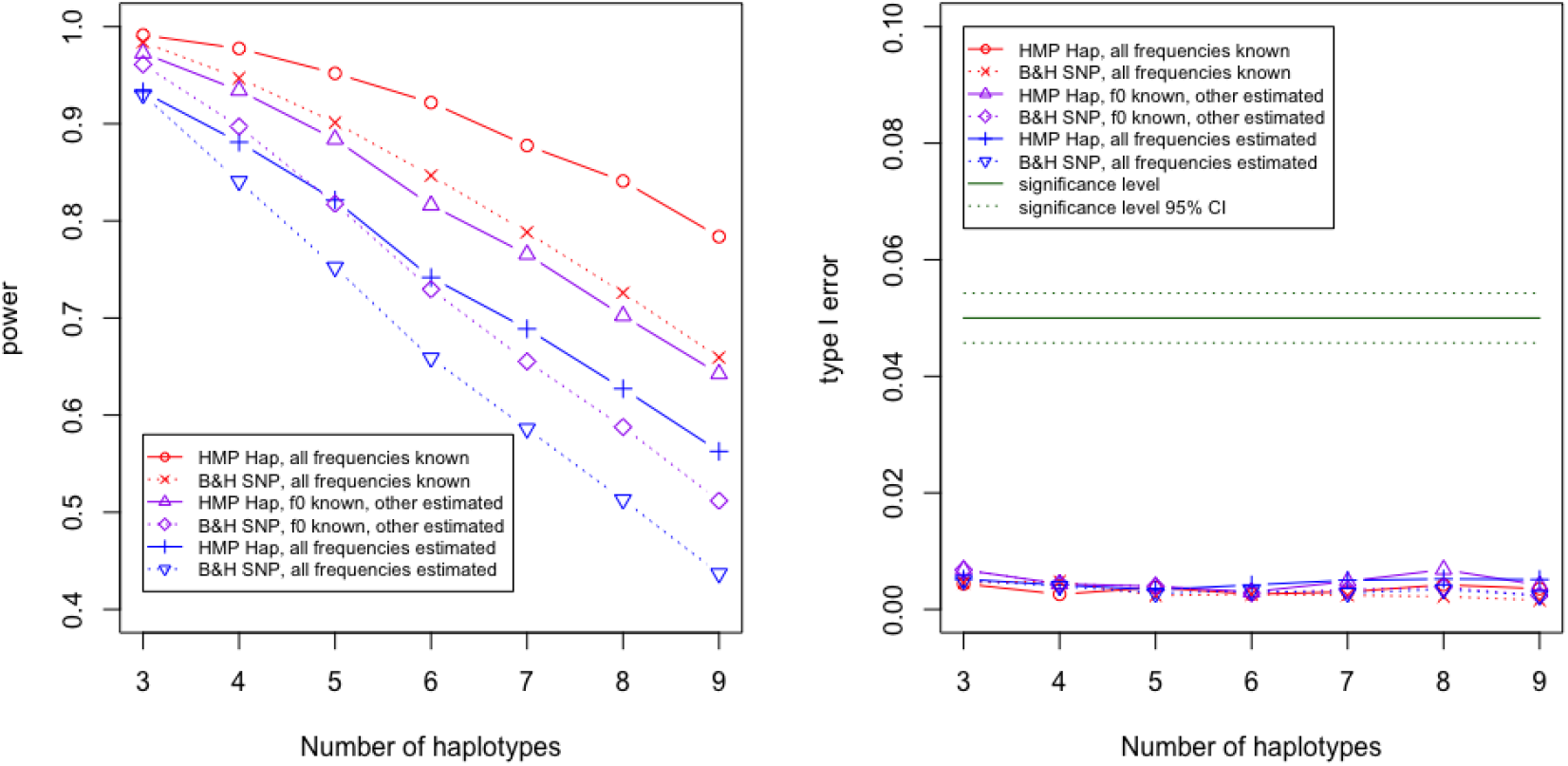
Results when changing the number of total haplotypes under 3 scenarios. Power and Type I error curves are plotted for both haplotype and SNP based test under various *N*_*Hap*_, with all other parameters default as outlined in Table 2 under 3 scenarios. Scenario 1: all frequencies are known. Scenario 2: starting frequencies are known, all other frequencies are estimated with a sample size of 80. Scenario 3: all frequencies are estimated with a sample size of 80.

#### S.7.2 Power and Type I error when both sampling and pool sequencing noise exists

**Figure S13:**
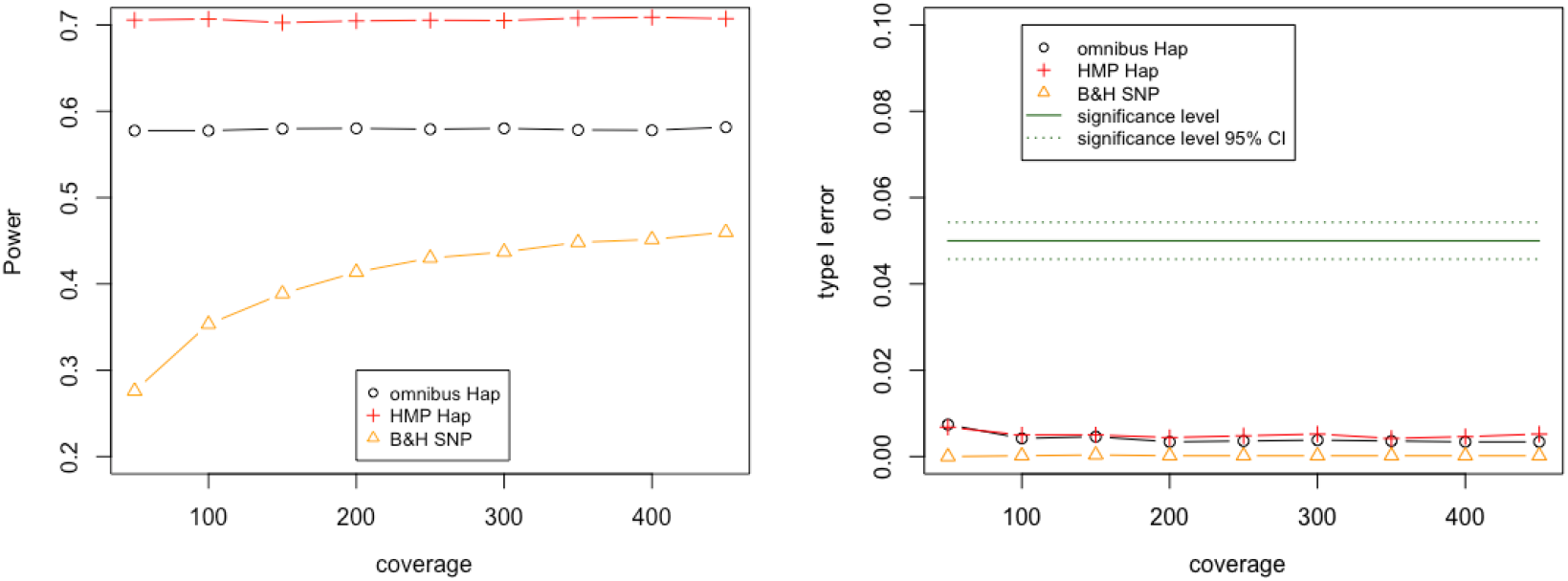
Results when changing the coverage. Power and type I error curves are plotted for haplotype and SNP based test under various sampling size. All frequencies are estimated with a sample size of 500, with other parameters default as outlined in Table 2.

### S.8 Testing for the number of selected haplotypes, unconditional on presence of selection

**Figure S14:**
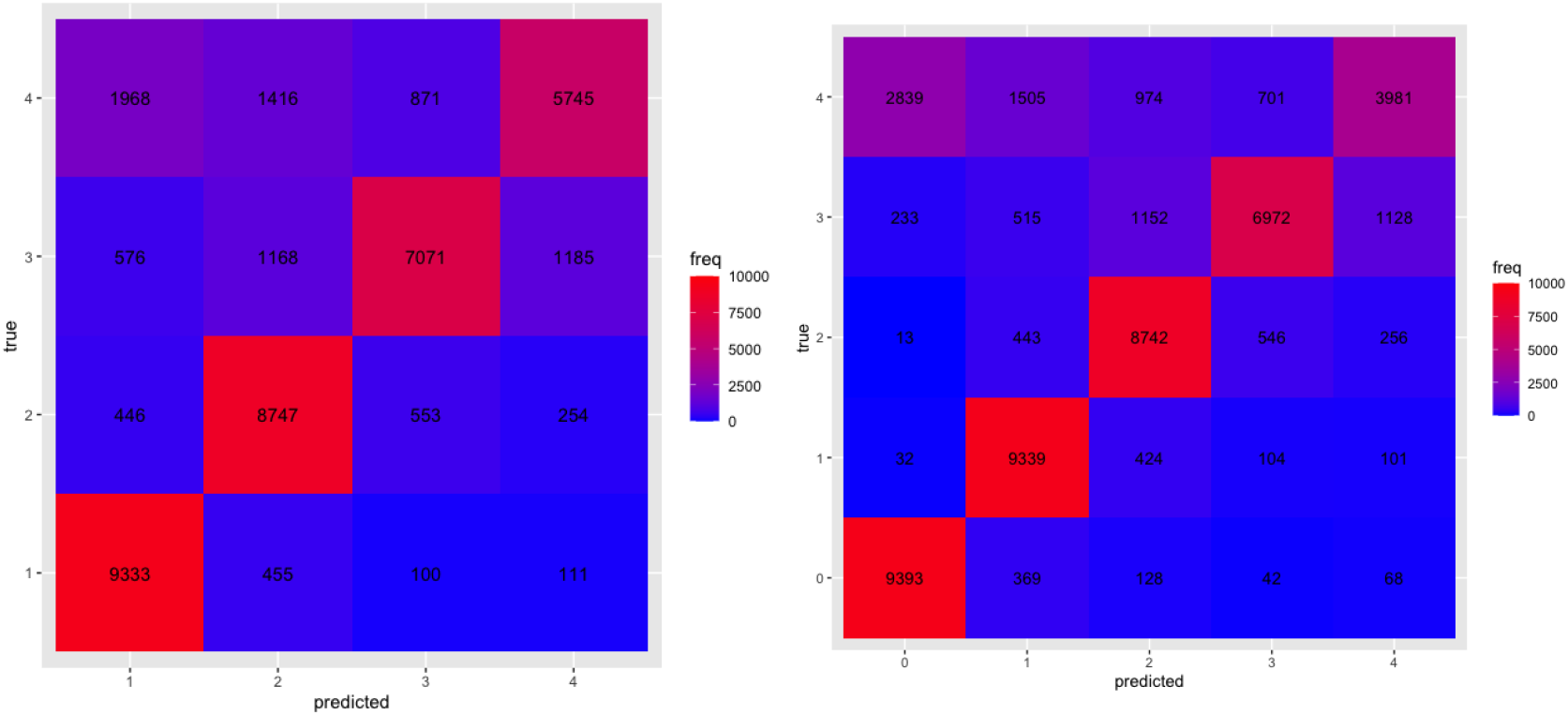
Result showing the predictability of haplotype based iterative testing. Omnibus is used for multiple testing correction under all scenarios. The heat map is simulated using 5 founder haplotypes and a selection strength of *s* = 0.05, 1 replicate population, with all other parameters default as outlined in Table 2. In the left panel, the test is conditional on the presence of selection. In the right panel, the test is unconditional.

### S.9 Testing for the number of selected haplotypes, using HMP as p-value combination method

**Figure S15:**
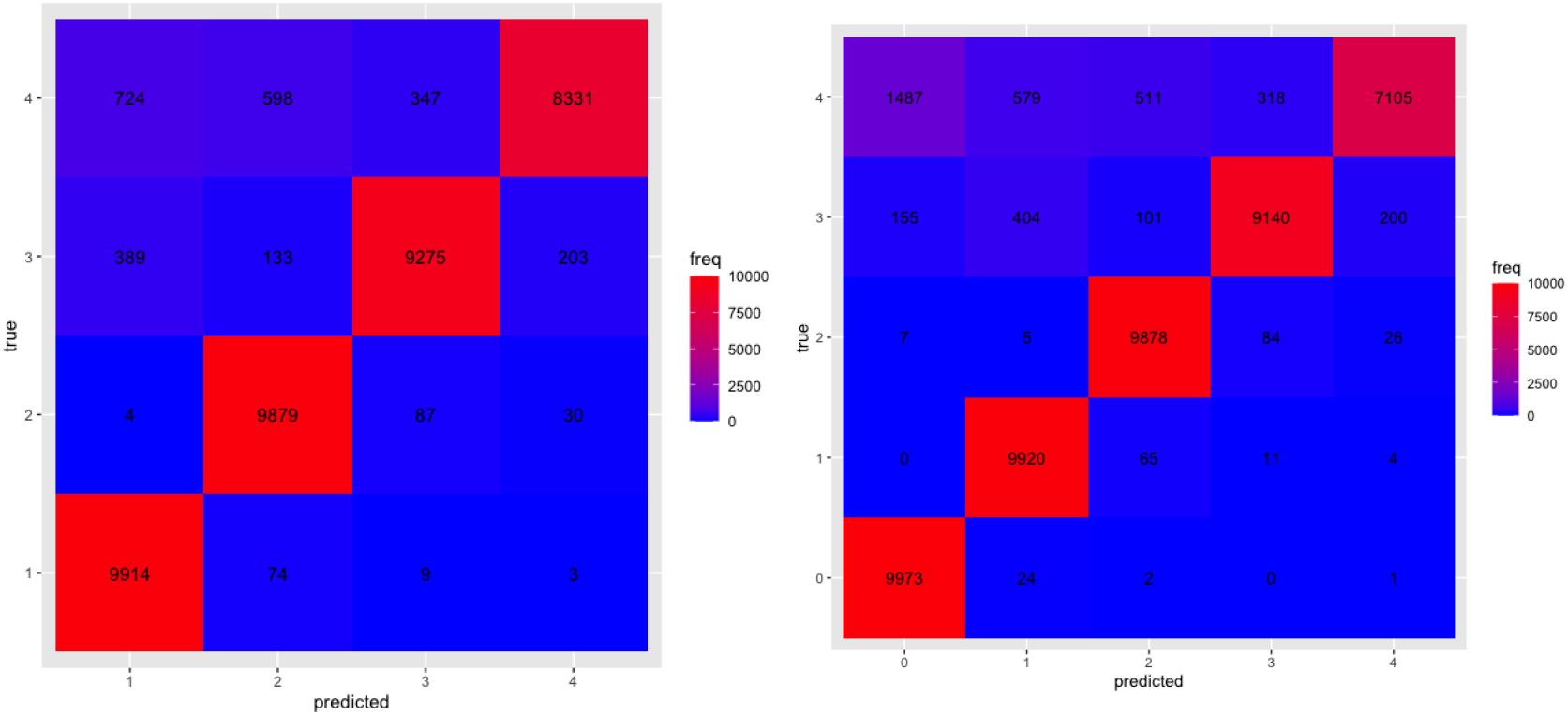
Results showing the prediction accuracy of haplotype based iterative testing. The HMP method is used for multiple testing correction in all scenarios. The heat map has been obtained using 5 founder haplotypes and a selection strength of *s* = 0.05, with all other parameters default as outlined in Table 2. In the left panel, the test is conditional on the presence of selection. In the right panel, the test is unconditional.

**Figure S16:**
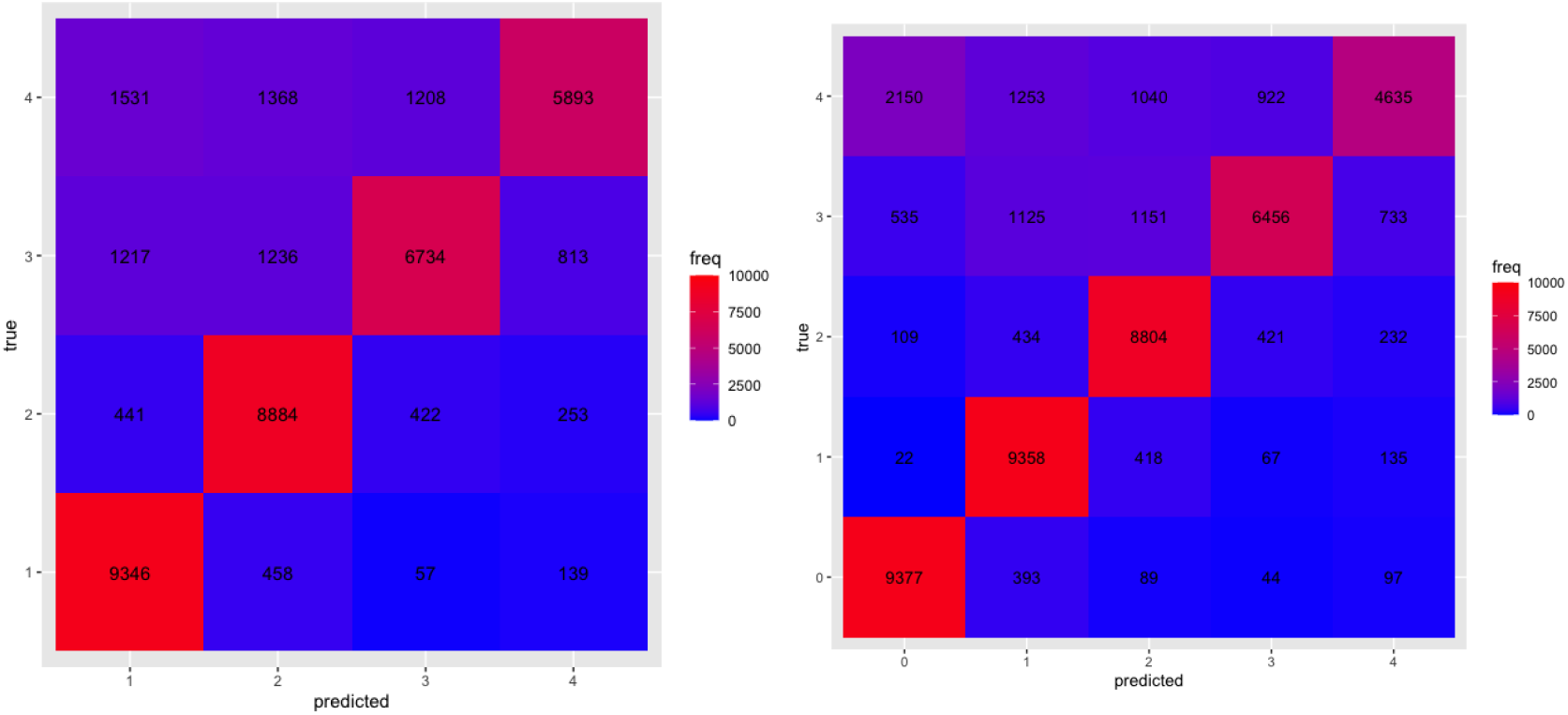
Result showing the predictability of haplotype based iterative testing. HMP is used for multiple testing correction under all scenarios. The heat map is simulated using 5 founder haplotypes and a selection strength of *s* = 0.05, 1 replicate population, with all other parameters default as outlined in Table 2. In the left panel, the test is conditional on the presence of selection. In the right panel, the test is unconditional.

### S.10 Type I error of testing for the number of selected haplotypes under low selective strength

**Figure S17:**
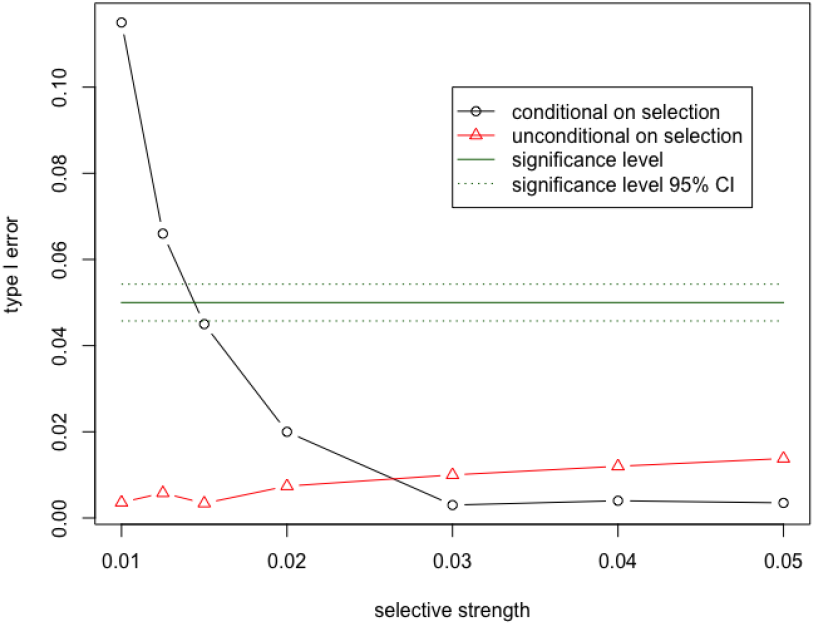
Type I error plot when testing for 1 vs more than 1 selected haplotypes. The type I errors were computed either conditional or unconditional on the rejection of the initial test for selection. The omnibus test is used for multiple testing correction. Data is simulated using 5 founder haplotypes and different selection coefficients *s*, with all other parameters as outlined in Table 2.

### S.11 Power of pairwise tests

**Figure S18:**
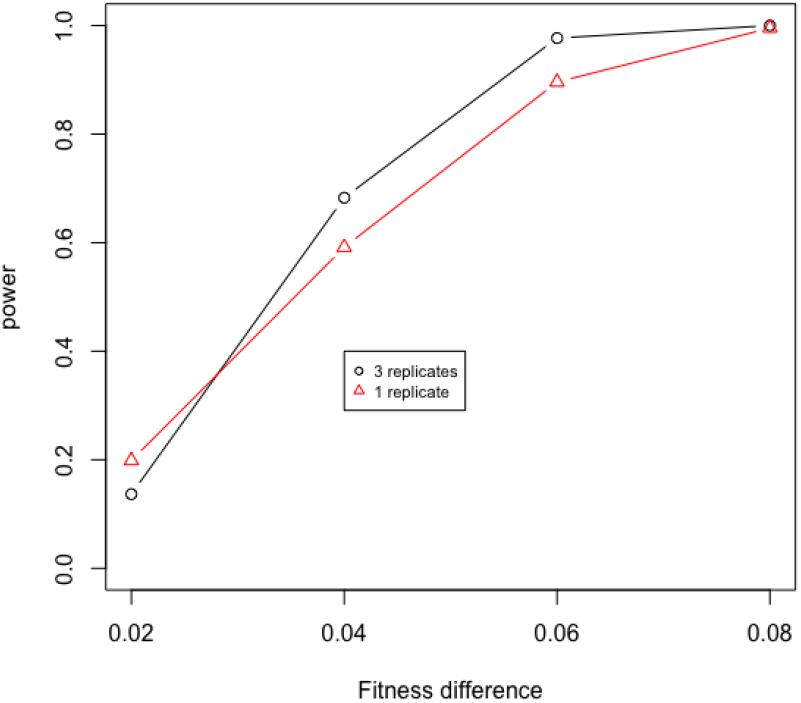
Results for pairwise tests when all haplotypes have different fitness. B&H is used for multiple testing correction. Data is simulated using 5 founder haplotypes, each having a fitness of (1,1.02,1.04,1.06,1.08), 1 or 3 replicates, with all other parameters default as outlined in Table 2.

The availability of replicate populations leads to more power for most scenarios. However it under-performs when the fitness difference is only 0.02. We notice a similar issue with replicate populations when applying our haplotype based test on cases where selected haplotypes have a very small selective advantage against others, see Section S.12. This is due to the adapted CMH test being more conservative than the adapted chi-squared test. If the selective strength is high enough, the information provided by the additional replicates makes the CMH test more powerful despite its conservativeness. When there is little signal (fitness difference 0.02), it leads to the lower power seen in both mentioned figures.

Since all fitness values are different to each other in this scenario, it is not possible to analyse type I error. Thus, we simulated one additional scenario where we randomly set two of the fitness values to 1 at each iteration, and observed that type I error is under control.

### S.12 Comparing chi-squared and CMH test under low selective strength

In Section S.11 we observed that the pairwise test with the adapted CMH test has lower power than with the adapted chi-squared test at very low fitness differences, despite the increase in replicates. We show in Figure S19 that, this also apply more generally to non-pairwise test scenarios. Counter-intuitively, here we observe that at very low selective strength, increasing the number of replicates resulted in lower power when applying the adapted CMH test.

**Figure S19:**
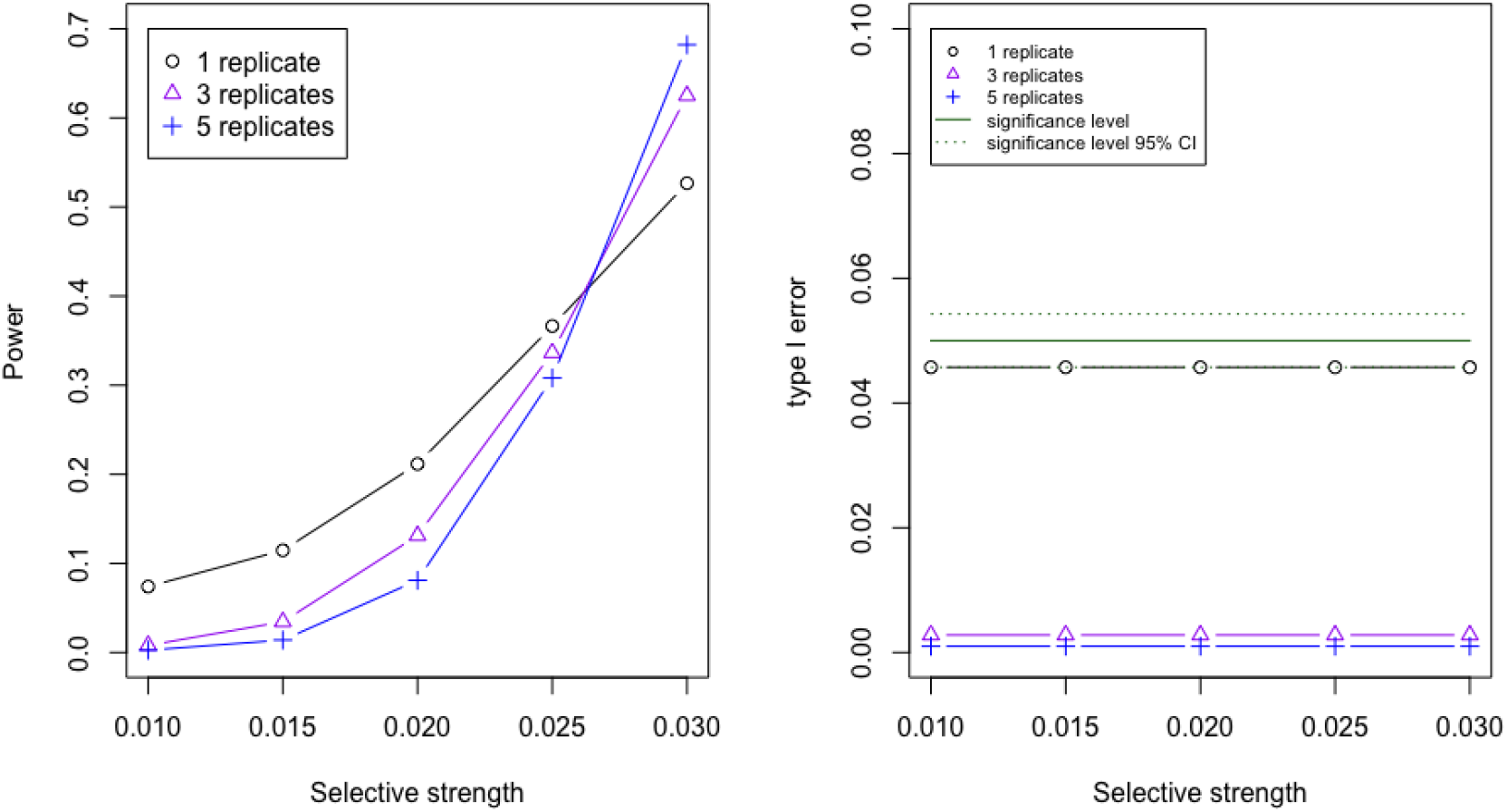
Results when changing selection strength and number of replicate populations. Power and type I error curves are plotted for haplotype based test using omnibus as p-value combination method under various *s* and *R*, with all other parameters default as outlined in Table 2.

This is caused by the CMH test being much more conservative than the adapted chi-squared test. In Figure S20 we show an additional scenario where we apply the adapted chi-squared test separately on each of the 3 replicates, and combine their results using a Benjamini & Hochberg multiple testing correction. Here we see that this method provides a significant increase in terms of power when compared to the scenario with one replicate, but also a much higher type I error when compared to the CMH test.

**Figure S20:**
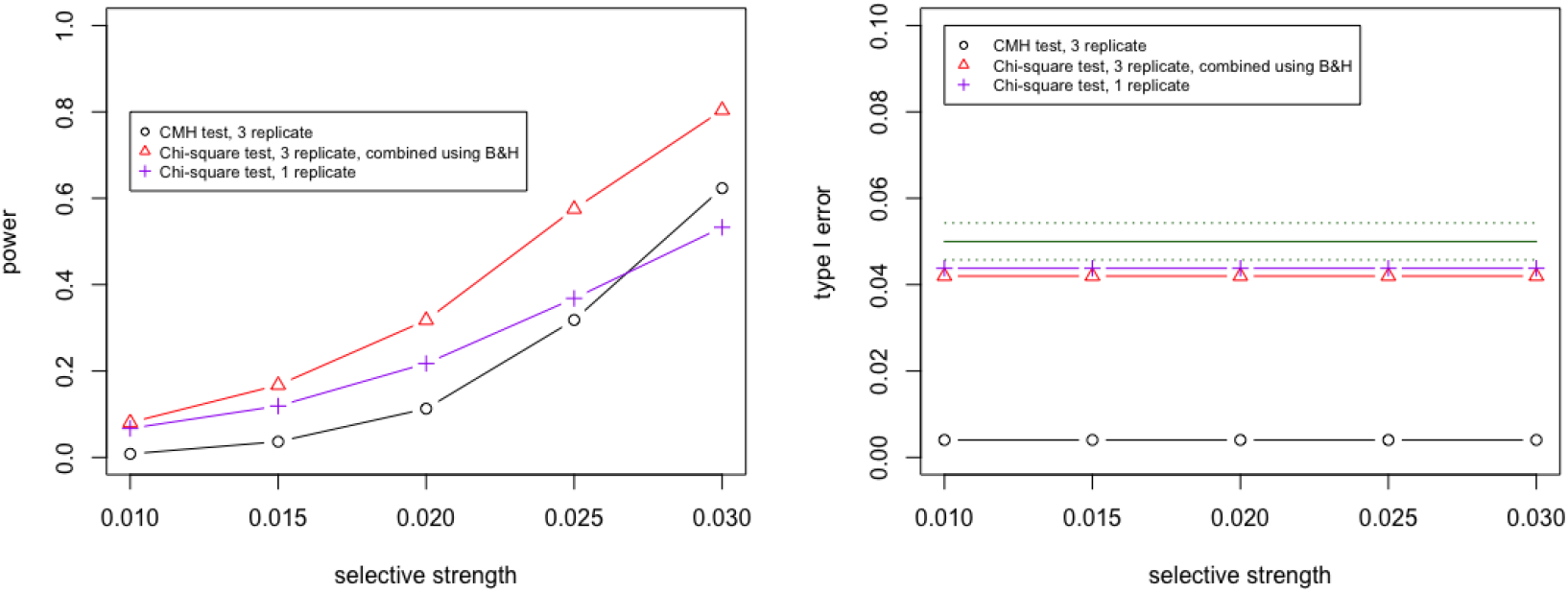
Results when changing selection strength, number of replicate, using 2 testing methods. Power and type I error curves are plotted for haplotype based test using omnibus as p-value combination method under various *s* and *R*, with all other parameters default as outlined in Table 2.

We also plotted ROC curves in Figure S21 and show that even at 0.02 selective strength, despite the low power and conservativeness, CMH test is still the best performing test overall for haplotype frequencies, in terms of AUC.

**Figure S21:**
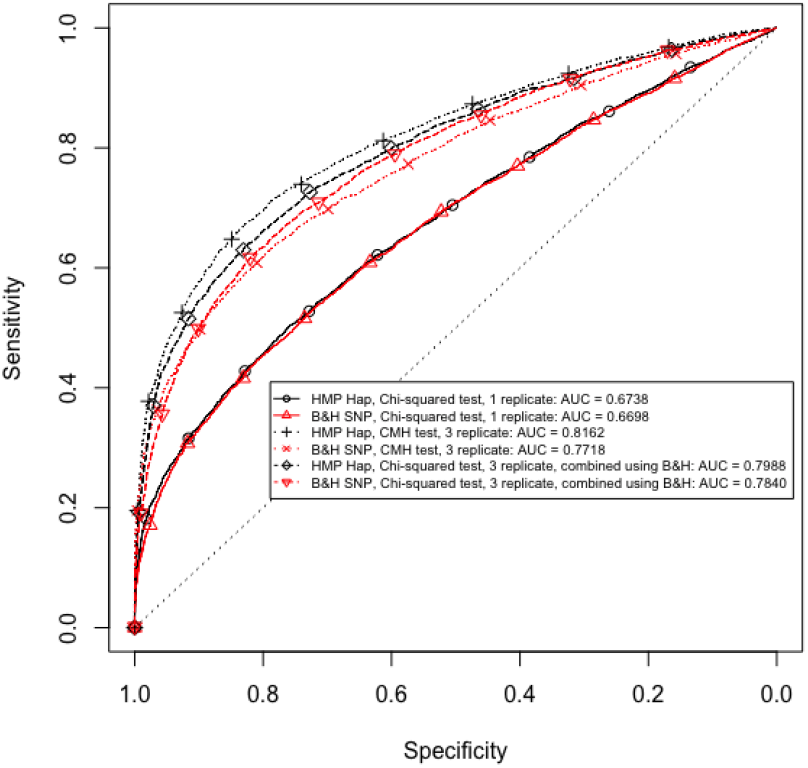
Results when changing the number of replicate and testing methods. ROC curves of haplotype and SNP based test when changing the testing method and number of replicates. The selective strength is set to be 0.02, with other parameters default as outlined in Table 2.

### S.13 ROC curves comparing haplotype combination methods when there is 100 haplotypes

**Figure S22:**
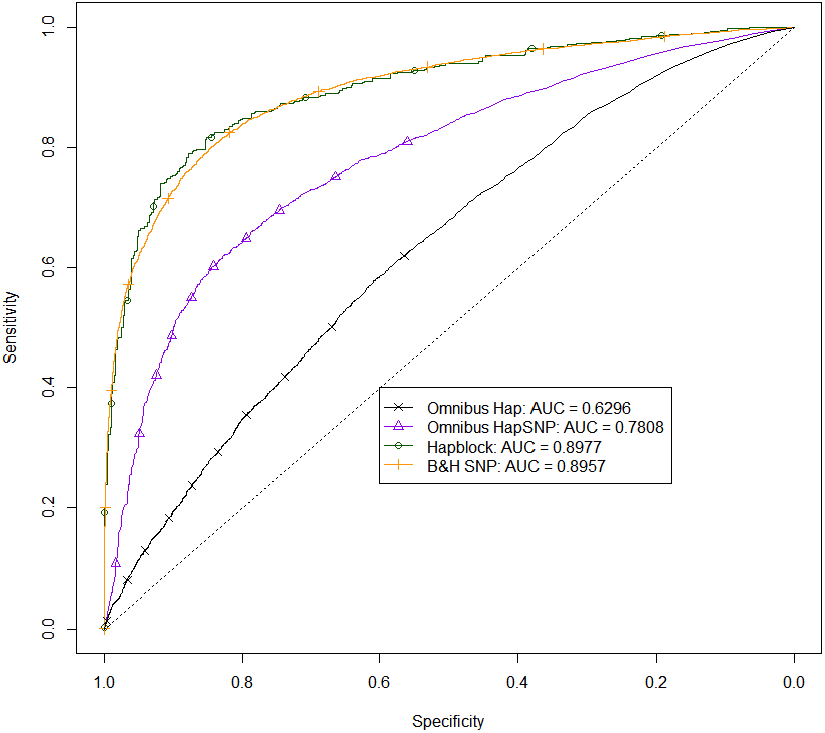
Results when there is 100 haplotypes. ROC curves of haplotype, HapSNP, hapblock and SNP based tests are shown for 100 total haplotypes, 50 selected haplotypes and no replicate population with all other parameters default as outlined in Table 2. Hapblock refers to the method of using haplotype blocks to reduce haplotype number outlined in Section 2.4.

### S.14 AUC comparing haplotype combination methods when changing the number of haplotypes

**Figure S23:**
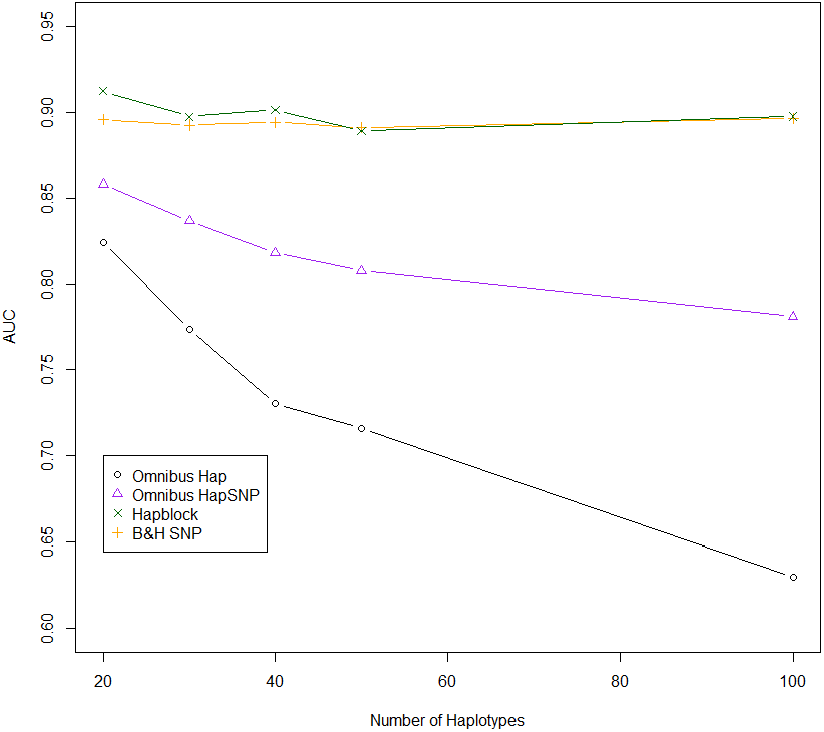
Results when changing the number of total haplotypes. AUC of the ROC curves are plotted for haplotype, HapSNP, Hapblock and SNP based test under various *N*_*Hap*_. Here, the number of selected haplotypes is set to be half of the total haplotype number *h*_*Sel*_ = *N*_*Hap*_*/*2, no replicate population, with all other parameters default as outlined in Table 2.

### S.15 Haplotype combination methods when only 1 selected haplotype exists

**Figure S24:**
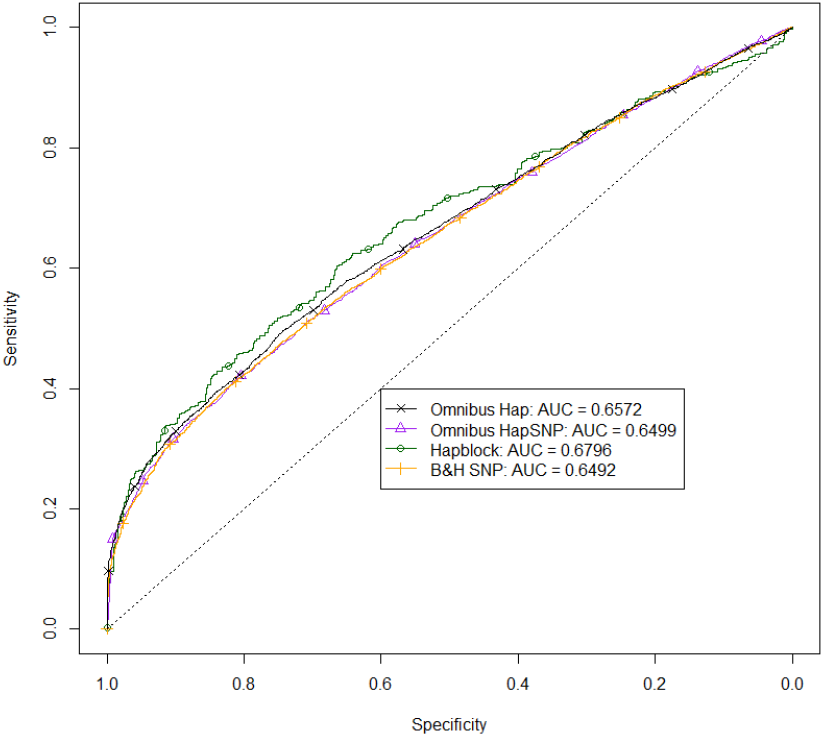
ROC curves when there are 30 haplotypes, one of them selected. Haplotype, HapSNP, Hapblock and SNP based tests are considered for 30 haplotypes, 1 selected haplotype and no replicate population, with all other parameters default as outlined in Table 2.

**Figure S25:**
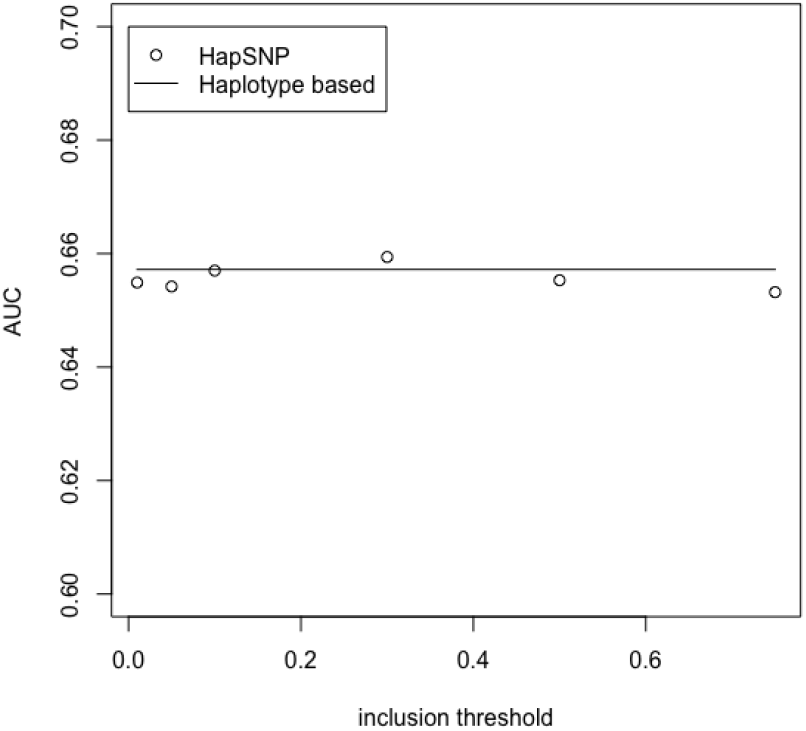
AUC when changing the inclusion threshold of SNPs. AUC for the HapSNP test is plotted under various inclusion threshold, compared with the AUC with haplotype based test, under parameters identical to Figure S24.

### S.16 Justification of not using cycle 12 in Section 4

We notice for population 4k an exceptionally high percentage of SNPs either fixed or lost at cycle 12, as shown in the table below:

**Table S1:**
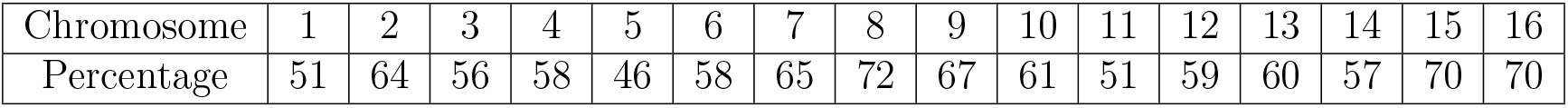
Percentage of SNPs either fixed or lost at last time point

This biases the SNP based test, as the fixation or loss of an allele can happen any time between cycle 6 and cycle 12, which would be roughly 105 generations. As no information is available between the cycles, we will not be able to obtain an accurate result using these data with the considered methods.

Moreover, the amount of average absolute frequency change between the first and second time point is much smaller than the frequency change between the second and third time point:

**Table S2:** Average absolute frequency change at each chromosome, between different cycles

| Chromosomes<br>Cycles | 1 | 2 | 3 | 4 | 5 | 6 | 7 | 8 | 9 | 10 | 11 | 12 | 13 | 14 | 15 | 16 |
| --- | --- | --- | --- | --- | --- | --- | --- | --- | --- | --- | --- | --- | --- | --- | --- | --- |
| 0 ~ 6 | 0.13 | 0.16 | 0.07 | 0.17 | 0.08 | 0.24 | 0.21 | 0.14 | 0.21 | 0.11 | 0.12 | 0.10 | 0.12 | 0.11 | 0.21 | 0.12 |
| 6 ~ 12 | 0.35 | 0.48 | 0.24 | 0.37 | 0.32 | 0.30 | 0.35 | 0.38 | 0.24 | 0.31 | 0.36 | 0.38 | 0.40 | 0.33 | 0.35 | 0.27 |

This might be explained by a shrinking population size, and thus much higher drift at latter cycles. Indeed, using the *N*_*e*_ estimates proposed by [36], we observe a much lower median *N*_*e*_ estimate between the latter cycles, which make the frequency changes noisier. It becomes therefore harder to separate selection and drift:

**Table S3:**
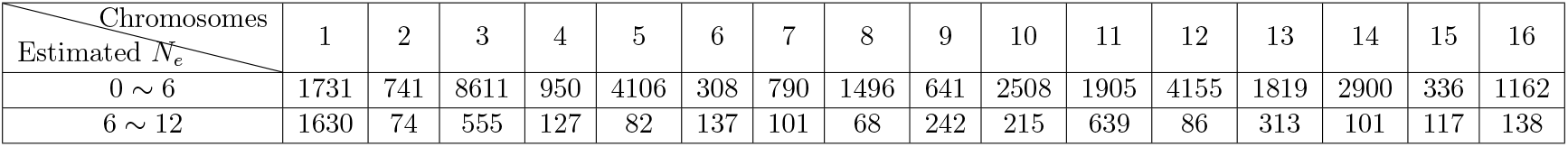
Median of estimated *N*_*e*_ using allele frequencies between different cycles

| Chromosomes<br>Estimated $N_e$ | 1 | 2 | 3 | 4 | 5 | 6 | 7 | 8 | 9 | 10 | 11 | 12 | 13 | 14 | 15 | 16 |
| --- | --- | --- | --- | --- | --- | --- | --- | --- | --- | --- | --- | --- | --- | --- | --- | --- |
| 0 ~ 6 | 1731 | 741 | 8611 | 950 | 4106 | 308 | 790 | 1496 | 641 | 2508 | 1905 | 4155 | 1819 | 2900 | 336 | 1162 |
| 6 ~ 12 | 1630 | 74 | 555 | 127 | 82 | 137 | 101 | 68 | 242 | 215 | 639 | 86 | 313 | 101 | 117 | 138 |

### S.17 Number of windows where selection is detected in real data application

**Table S4:** Absolute number of windows each method is able to detect as selected for each chromosome.

| Chromosomes | Total | 1 | 2 | 3 | 4 | 5 | 6 | 7 | 8 | 9 | 10 | 11 | 12 | 13 | 14 | 15 | 16 |
| --- | --- | --- | --- | --- | --- | --- | --- | --- | --- | --- | --- | --- | --- | --- | --- | --- | --- |
| Hap based | 1522 | 24 | 39 | 62 | 209 | 182 | 23 | 42 | 15 | 39 | 193 | 136 | 80 | 100 | 129 | 115 | 134 |
| SNP based | 802 | 35 | 17 | 85 | 147 | 0 | 0 | 157 | 94 | 0 | 40 | 0 | 51 | 54 | 8 | 51 | 63 |
| Total window number | 11482 | 186 | 788 | 294 | 1496 | 550 | 240 | 1046 | 534 | 396 | 687 | 633 | 1038 | 905 | 751 | 1036 | 902 |

